# Readiness potential in human functional magnetic resonance imaging motor task data

**DOI:** 10.1101/2025.01.31.635910

**Authors:** Marika Strindberg, Yu Tian, Peter Fransson

## Abstract

The readiness potential (RP), also known as the Bereichtschaftspotential, is commonly observed in EEG recordings as a slow build-up of negative electrical potential prior to self-directed motor movements (Kornhuber et al. 1965). In this study, we analyzed motor task-based functional magnetic resonance (fMRI) data acquired from 262 human participants, focusing on the degree of signal phase synchronization across different frequency spans and brain regions. A new method that clusters brain regions based on instantaneous phase allowed for time resolved estimation of cross-subject phase synchronization. We show that during rest periods that precede movement, an fMRI equivalent of the RP is gradually established in a network that encompasses the bilateral supplementary motor area and parts of the cingulo-opercular network, recently described as the brain’s Action Mode network (Dosenbach et al. 2025). Importantly, an anticipatory negative shift in the fMRI signal was observed in both single subjects and single epochs. Tongue movement elicited strong synchronization between the orbitofrontal, ventromedial, and temporal pole cortices. Our findings suggest a link between fMRI and electrophysiological recordings of anticipatory motor events in the brain. The method presented here grades brain synchronization along an in-phase and anti-phase continuum and has applications in clinical settings, as well as for cognitive brain mapping that goes beyond anticipation.

**Significance statement:** The readiness potential (RP) is observed in EEG recordings as a slow build-up of negative electrical potential in the supplementary motor area (SMA) starting up to two seconds before self-initiated motor movements. It is believed to play a causal role in motor preparation. Given that it is only detectable by averaging 30-40 epochs, its existence as a real signal has been challenged by new modeling approaches which suggests the early RP-component is merely a summation of stochastic noise. In this study, we demonstrate that the equivalent of the RP is present in functional magnetic resonance imaging (fMRI) data in a distributed network known as the “action mode network”. The RP-equivalent is present in single subjects and single epochs.

## Introduction

The readiness potential (RP) is characterized by a gradual increase in negative potential, observable in time-locked averages of neurophysiological recordings of the brain preceding self-initiated motor movements^1–3^. The RP comprises early and late components. The early component commences up to 2 s prior to motor movement and predominantly reflects bilateral activity in the supplementary motor area (SMA), pre-SMA, and anterior mid-cingulate cortex. The late component initiates approximately 400 ms before movement onset and is lateralized to the hemisphere opposite the motor movement, corresponding to activity in the motor cortex ^4^. By contrast, self-reports and empirical evidence indicate that the intention to act typically arises 200 ms before a motor action ^5^.

Since its initial observation in the 1960s^3^, the readiness potential (RP) has been extensively investigated^1,6–8^. However, its functional significance and existence as a genuine phenomenon in the brain remain debatable. The discrepancy in timing between self-reports of the decision to act and the onset of the RP has even been employed as an argument against the concept of free will^9^. The RP is most readily observed prior to self-initiated movement, wherein participants are instructed to move when they feel the urge to do so. This contrasts with cued movements, in which participants moved only after receiving a cue. Although the RP is also observed in the latter context, it manifests with reduced amplitude^6^. Common to these scenarios is the perceived imperative to move, with the distinction that timing is less constrained in the self-paced scenario.

The classical interpretation posits that the readiness potential (RP) plays a causal role in motor preparation. However, RP has also been observed in instances where participants decide to move but do not execute the movement^6^. This challenges the strict classical interpretation and suggests that RP may reflect a more general decision-making process. Consequently, it remains uncertain whether the RP is indeed distinct from other negative potentials observed prior to cognitive tasks in cued paradigms that do not necessarily involve a motor component, such as the contingent negative variation (CNV) and stimulus-preceding negativity (SPN)^2^. The CNV and SPN are generally considered to reflect anticipation and attention, engaging brain areas essential for upcoming tasks ^2^.

A more recent explanatory framework, supported by modelling approaches and empirical data, has provided a radical revaluation of the RP ^10^. The stochastic accumulator model proposes that the RP may not be a genuine signal but rather an artifact resulting from averaging time-locked stochastic noise. According to this model, the decision to move occurs when the accumulation of noise reaches a certain (undefined) threshold. Thus, averaging the epochs of movement introduces a selection bias where the pre-threshold data are inherently lower. Consequently, this model suggests that the post-threshold or late RP component reflects a decision process, whereas the pre-threshold build-up represents stochastic noise.

Recent empirical data suggest that slow cortical potentials (SCP) could account for the RP phenomena^11^. SCPs are slow fluctuations (0.01–1 Hz) in the cortical potential in neurophysiological recordings that are spontaneously present^6,11^. There is strong evidence of a correlation between SCPs and slow fluctuations in BOLD fMRI ^12^. A negative shift in SCPs has been linked to increased responsiveness and cognitive performance ^11^. Experimental findings indicate that both the subjective urge to move and the likelihood of movement are greater during a negative shift than during a positive shift in SCP. The SCP sampling hypothesis posits that the RP represents the subjective urge to move ^6,11^. From this perspective, the early component of the RP is the summation of the predominantly negative shifts in the SCP preceding movement. It has also been demonstrated that it is possible to modulate the SCP by intention and delay movement onset, even when a strong negative shift, indicative of an urge, is present^11,13^.

In summary, three principal explanatory frameworks for the RP have been proposed: 1) a direct causal role in preparing motor action, rendering it necessary for motor movement; 2) a result of stochastic fluctuations that trigger a decision to move upon reaching a certain threshold, thereby rendering the early component of the RP an artifact; and 3) a reflection of the likelihood of ongoing fluctuations in cortical potential, where negative shifts create an urge to move.

A central issue contributing to the varied interpretations of the readiness potential (RP) is the low signal-to-noise ratio in electroencephalography (EEG), which requires averaging across 30-40 time-locked epochs or several electrodes to observe the RP^1,2,14^. A crucial step towards a more comprehensive understanding of the RP involves unequivocally determining whether the RP exists within single epochs or is merely an artifact of averaging.

In this study, we leverage the enhanced signal-to-noise ratio and spatial precision of functional magnetic resonance imaging (fMRI) to address this question. We employed a new time-resolved data analysis method using a substantial sample of the Human

Connectome Project dataset^15,16^, focusing on fMRI motor task data involving hand, foot, and tongue movements across two distinct BOLD signal frequency domains, 0.066-0.189 Hz and 0.030-0.010 Hz. Our findings show that within the lower frequency domain, the equivalent of the readiness potential (RP) emerges towards the end of fixation periods between task blocks. It is observed bilaterally in the supplementary motor area (SMA) and most other regions of the cingulo-opercular action mode networks (CO-AMN). Notably, we demonstrate that this gradual and increasingly negative shift in the blood-oxygen-level-dependent (BOLD) signal is readily observable in single epochs and individual subjects.

## Results

### Intrinsic mode signal extraction and phase community assignment

Functional imaging data from the two motor tasks in the fMRI HCP dataset (282 participants) were parcellated using a standardized parcellation scheme^17^ (results from the final run are presented here, with differences between runs noted when relevant). Bandpass-filtered intrinsic mode signals from each parcel were extracted using the complete ensemble empirical mode decomposition (CEEMD) algorithm^18^ (Suppl. Fig. S1). The analysis was restricted to two intrinsic modes that represented signal changes in a lower (0.010-0.030 Hz) and upper (0.066-0.189 Hz) frequency band. Instantaneous phase was computed at each time point and intrinsic mode time series. A phase-clustering algorithm was developed to form communities of parcels that consistently spanned a limited phase range across time and subjects (Fig. 1). The algorithm was designed such that any pairwise relationship between parcels within the same community would have an instantaneous phase difference that did not exceed a predefined threshold. The results were six major and two minor communities for both frequency bands (referred to as COMs). The major COMs were similar in size, with each covering a limited section of the unit circle (Fig. 1A), with some overlaps (Fig. 1B). Consequently, each community consistently represented approximately the same instantaneous phase, that is, sections of positive and negative shifts (deflections) of the time series (Figs. 1B, C). This provided a simple way to compare time series across parcels, time points, and subjects. The major COMs were numbered sequentially (COM1 – COM6) as they appear in the unit circle in Fig. 1A. To assess the statistical significance of the results, the time series were phase-shuffled, and the same analysis was applied to the phase-shuffled data.

**Fig. 1.**
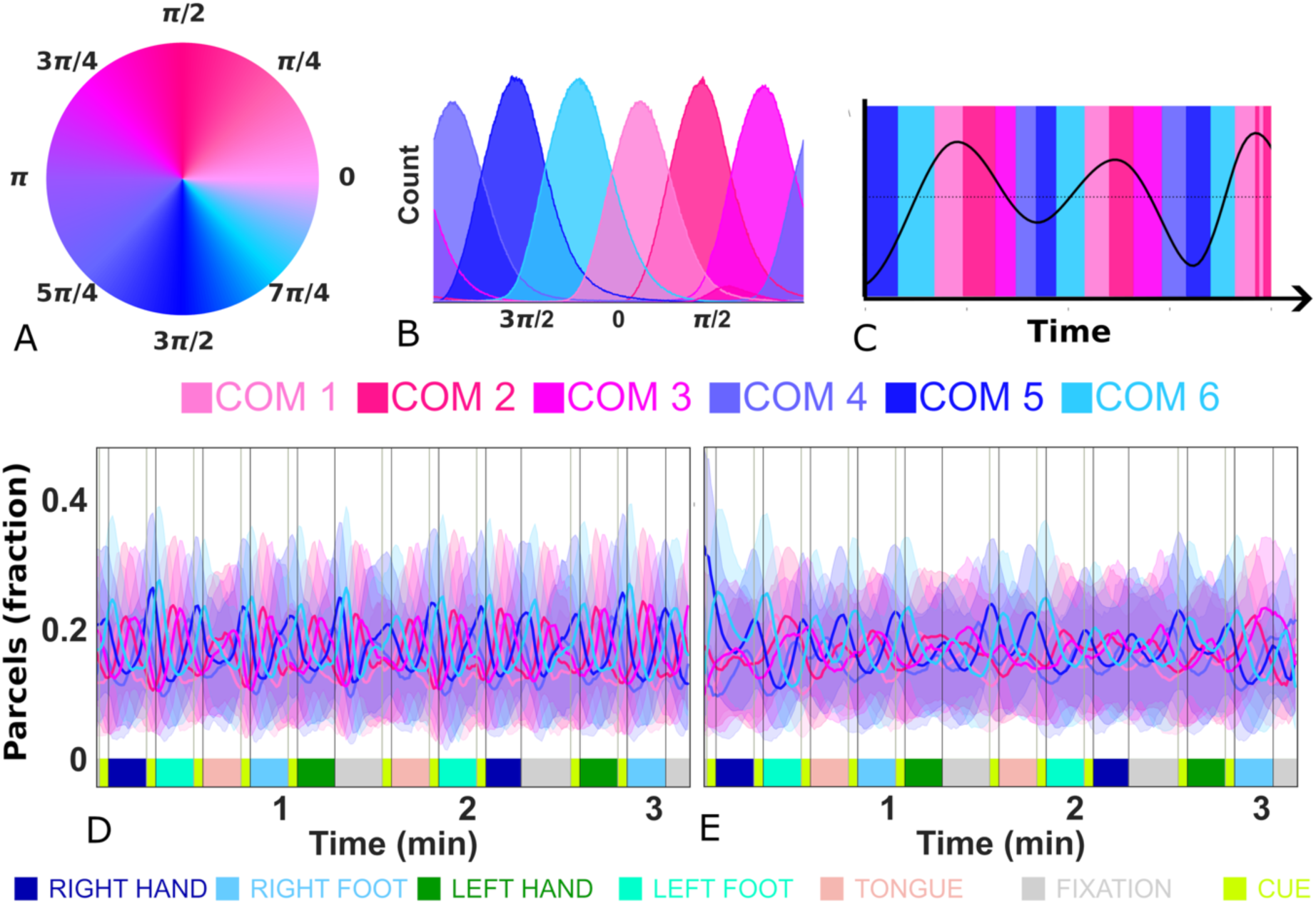
Community assignment based on the instantaneous phase of motor task fMRI data. **A)** Color wheel in which each instantaneous signal phase corresponds to a color. **B)** Distribution of the instantaneous phase (radians) across subjects, parcels, and time points) per community (plot is cut at *π*). For color coding, the median value of the phase distribution in each community was used based on (A). **C)** Correspondence between an intrinsic mode signal time course (lower frequency band) and community membership (COM1-C) for a single subject and parcel. For example, COM5 approximately represents the maximum points of negative deffection (or signal shift) over time. The average fraction of parcels (out of 23C) for each community across time was computed for the upper (0.0CC-0.18S Hz) (**D**) and lower (0.030-0.010 Hz) frequency bands (**E**).

### Two types of task-locked synchronization

Our analysis identified two types of motor task-locked synchronizations across subjects for both frequency bands. The first type involved regular fluctuations in the average COM size and was observed for all the COMs. These fluctuations were time-locked to the paradigm and were independent of the specific motor task performed. The oscillatory peaks in size corresponded to the switching order of the respective COMs, reflecting the progression of the signal phase (see Figs. 1D-E). In both frequency bands, the size of COM5, which aligned approximately with the peaks of the negative shifts in the time series (Figs. 1B-C) peaked at each cue (and at the time point where the cue would have been for the fixation blocks). The remaining five major COMs exhibited peaks in size in progressive order, thereby completing a phase cycle within the period of a task block (cue + task) in both the upper and lower frequency bands.

The second type comprised brain-network-specific patterns of phase synchronization across subjects. Cross-subject phase synchronization was defined for each parcel as the percentage of subjects sharing the same community assignment at a given time, thus providing a measure of the degree to which the phase of each parcel time series was synchronized across subjects. For the upper frequency band (Fig. 2), peaks of synchronization were observed for the visual (VIS) and attention networks (ventral attention, VAN, and dorsal attention, DAN), suggesting engagement independent of the specificity of motor tasks and instead mirroring the structure and timing of the block design. To a lesser extent, this phenomenon was also observed in segments of the frontoparietal network (FPN), basal ganglia (BG), thalamus (Thal), and for a few parcels in the default mode network (DMN) (Fig. 2). Marked synchronization in COM5 aligned with the onset of the cues was followed by a sequence of phase progression delineated by peaks in synchronization in COM6, COM1, COM2, COM4, and COM4. Thus, a complete cycle of the signal phase was achieved within the duration of a single motor task block (i.e., 3s cue + 12s task). This corresponds to a distinct peak in the power spectrum at 0.066 Hz, representing the 15s block design (Suppl. Fig. S1). Most parcels in the visual network exhibited cross-subject desynchronization during the fixation blocks (Suppl. Fig. S2A-B), indicating that the phase of the parcel time series was no longer synchronized across subjects. This desynchronization phenomenon was also observed in a few parcels of the attention networks.

**Fig. 2.**
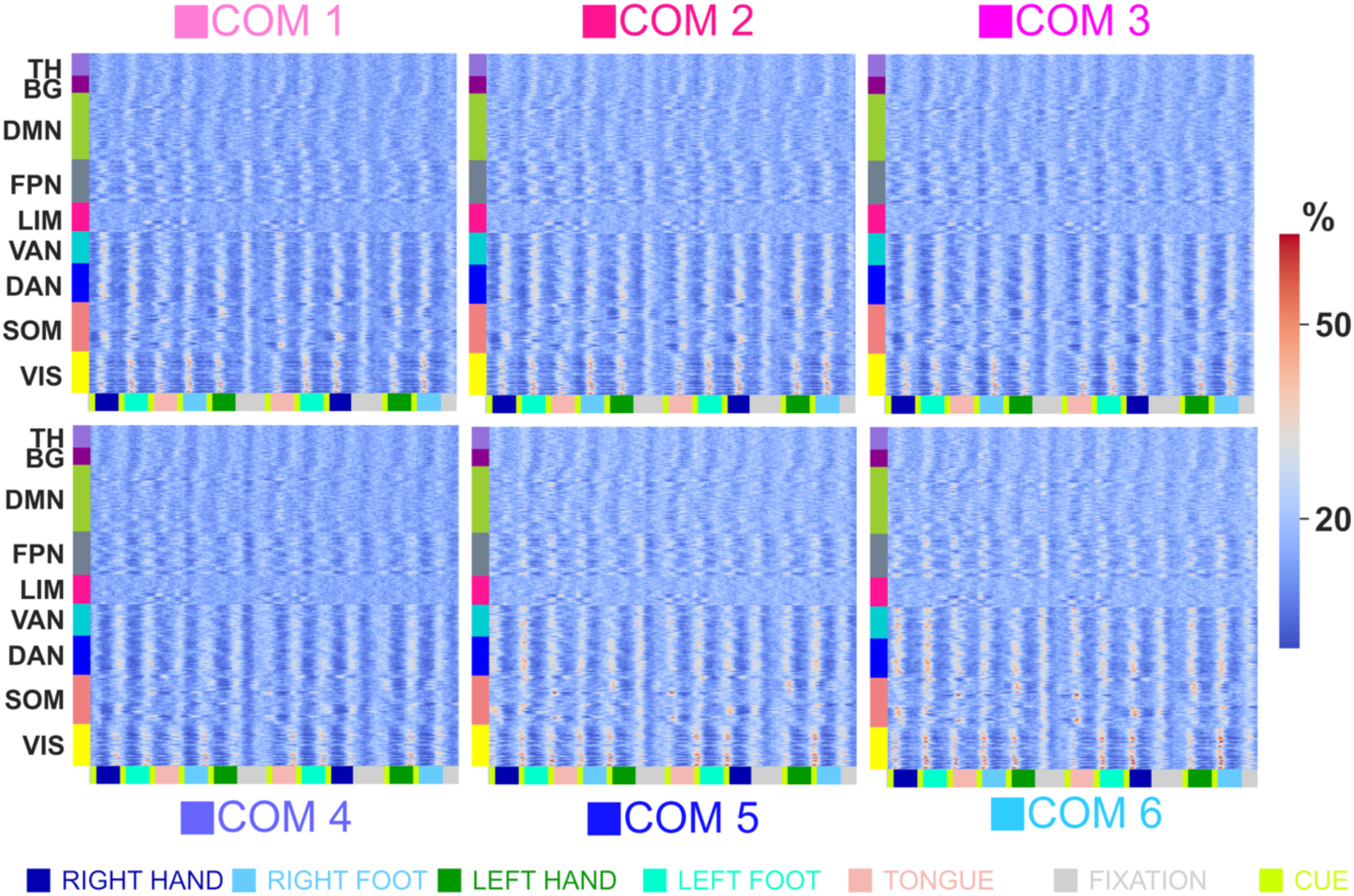
Network-specific cross-subject phase synchronization (n=282) in six communities (COM1-C) during an fMRI motor task (results shown here refer to signal changes in the upper frequency range of: 0.0CC-0.18S Hz). The color bar reffects the percentage of the total number of subjects (282 subjects) that were synchronized in each community at each point in time.

In the lower-frequency band (Fig. 3), COM membership patterns were unique to each of the five motor tasks (left/right foot, left/right hand, and tongue movement). Notably, specific subcohorts of parcels within the somatomotor network (SOM) were selectively synchronized, depending on the specific body part being moved (Fig. 3). Additionally, peaks in cross-subject phase synchronization were present during fixation periods in regions associated with anticipation and planning of motor movements, a result that will be elaborated later. While the block design-induced responses resulted in cycles of cross-subject phase synchronization that were completed within a task block for the upper frequency band, the period of task-related cross-subject phase synchronization in the lower frequency band persisted over approximately two and a half blocks (33 s) (Fig. 3, Suppl. Fig. S1).

**Fig. 3.**
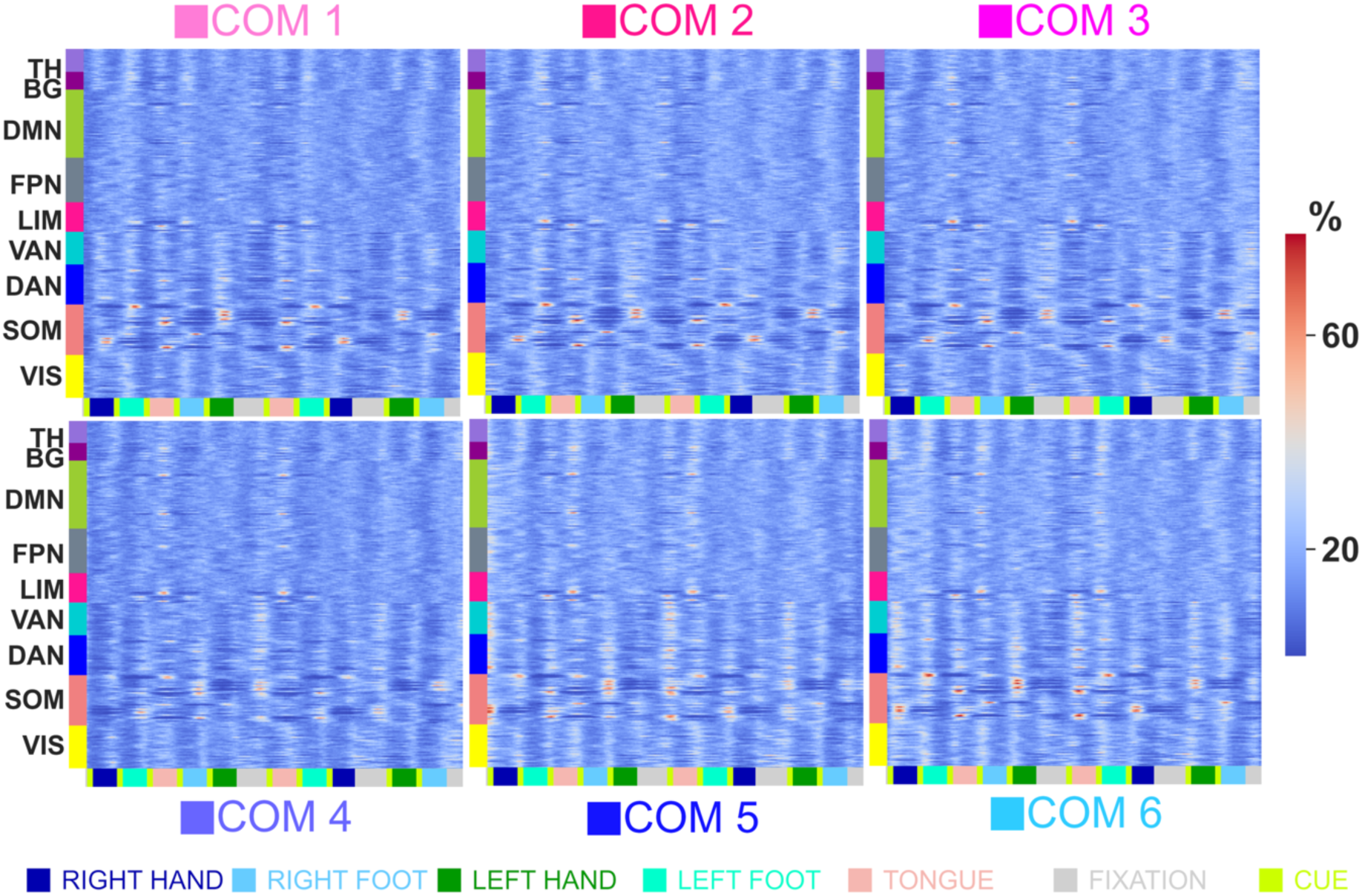
Network-specific cross-subject phase synchronization (n=282) in six communities (COM1-C) during an fMRI motor task (results shown here refer to signal changes in the lower-frequency range: 0.030-0.010 Hz). The color bar reffects the percentage of the total number of subjects (282 subjects) that were synchronized in each community at each point in time.

### Task-locked synchronization in the somatomotor network

Fig. 4A shows the intrinsic mode signals (lower frequency band) for five parcels in the motor cortex directly involved in the execution of hand, foot, and tongue movements. The corresponding progression of the cross-subject phase synchronization for each of the six communities is shown in Fig. 4B. Importantly, the cross-subject phase synchronization in Fig. 4B adds time-resolved information on the statistical significance of the averaged intrinsic mode signal in Fig. 4A in terms of the percentage of the total number of subjects aligned in phase. Statistical significance thresholds of cross-subject phase synchronization were calculated from the phase-shuffled data (Suppl. Fig. S3). There was a strong positive relationship between intrinsic mode signal amplitude and the maximum (across the six communities) cross-subject phase synchronization (Suppl. Fig. S4). The whole-brain maps shown in Fig. 4C depict the degree to which other brain areas were correlated in terms of synchronization (across subjects) with the five seed parcels. This shows that several regions beyond the five parcels in the motor cortex, such as the associated somatosensory cortex and neighboring dorsal regions in the dorsal attention network, were synchronized during motor task execution. Notably, the highest degree of cross-subject phase synchronization was observed in the hand somatosensory cortex. Tongue movements induced nearly equal-sized synchronizations in the associated motor regions in both hemispheres. It also induced significant synchronization with the orbitofrontal cortex and the temporal poles (Suppl. Fig. S7,8). Hand activation was unilateral, whereas motor parcels of the feet exhibited weak synchronization when the contralateral foot moved.

**Fig. 4.**
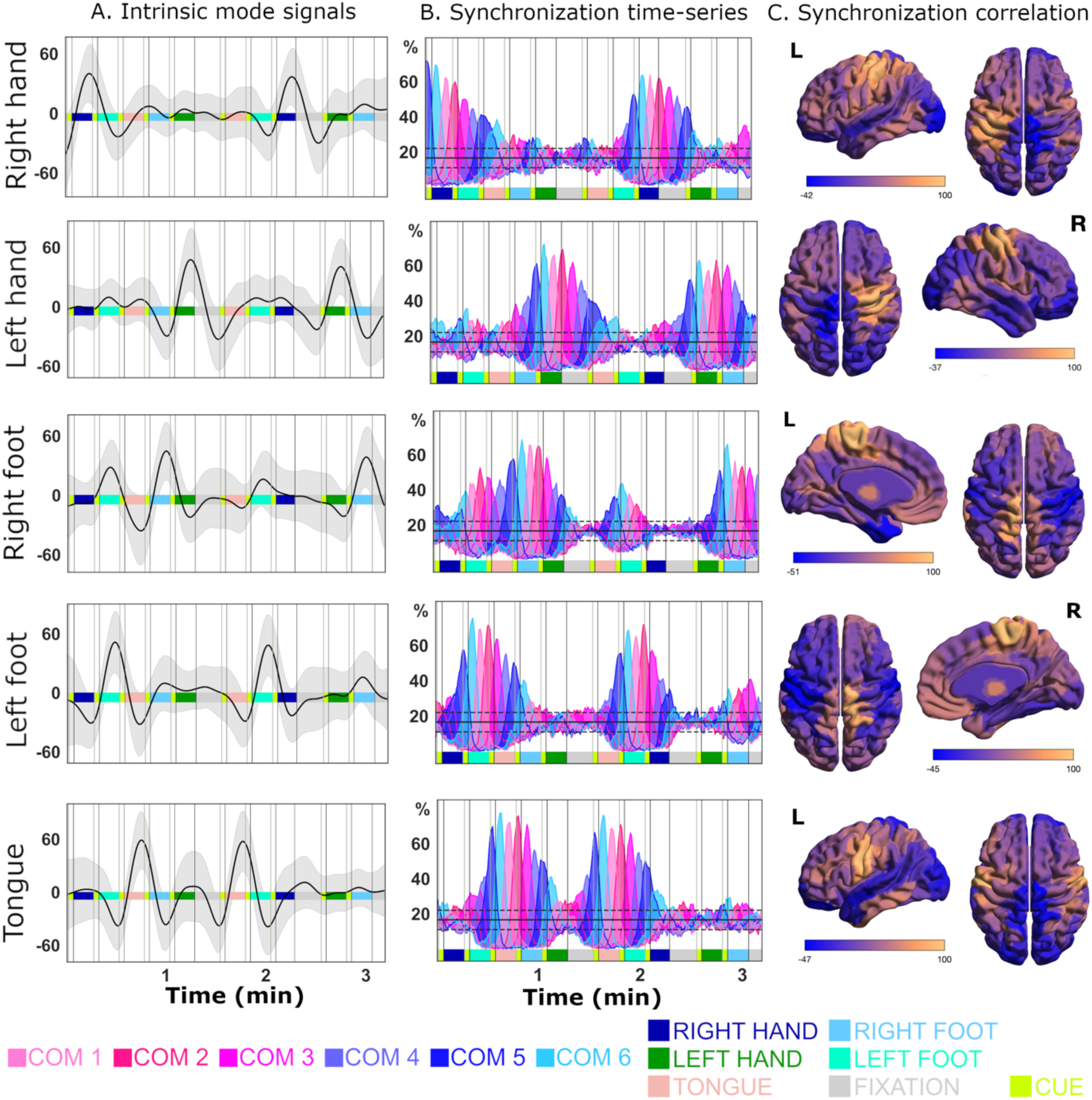
Averaged (n=282) intrinsic mode signals (lower frequency band, shaded area = ± 1 SD) **(A)** and the fraction of subjects that were phase-synchronized into COM1-C across time for the fMRI motor task **(B)** for five parcels in the motor cortex (SOM network) responsible for movement execution: hands (LH_SomMot_C, RH_SomMot_14), feet (LH_SomMot_12, RH_SomMot_14), and tongue (RH_SomMot_18). The correlation maps depicted in **(C)** show the average degree of correlation (Pearson’s r × 100) between the seed parcels in the motor cortex and the rest of the brain in terms of the percentage of synchronized subjects (correlation analysis performed separately for each COM and then averaged). The y-axis in (B) represents the fraction (in percentage) of subjects being synchronized. The horizontal black lines denote the mean degree of cross-subject synchronization (1C.C7%) for the phase-shu=led version of the data (dotted lines = ± 2 SD). Only the left hemisphere motor cortex representation of the tongue is shown (right side strongly correlated, Pearsons’ r = 0.S7). See Suppl. Fig. S6 for the averaged raw BOLD signals. The amygdala, hippocampus, and basal ganglia parcels are not shown here.

### Patterns of whole-brain task-locked synchronization

A whole-brain perspective of community phase synchronization during an fMRI motor task is shown in Fig. 5. The primary visual cortex (V1) and segments of the ventral visual stream, along with the primary somatosensory and motor areas of the tongue (more pronounced on the right side), demonstrated the highest degree of synchronization within the upper frequency band (Fig. 5). The somatosensory and motor regions were associated with the maximum degree of synchronization in the lower frequency band (Fig. 5B). We computed the average correlation between the parcel community synchronization time series (mean across the six main communities and all subjects) and sorted them according to network (Figs. 5C, D).

**Fig. 5.**
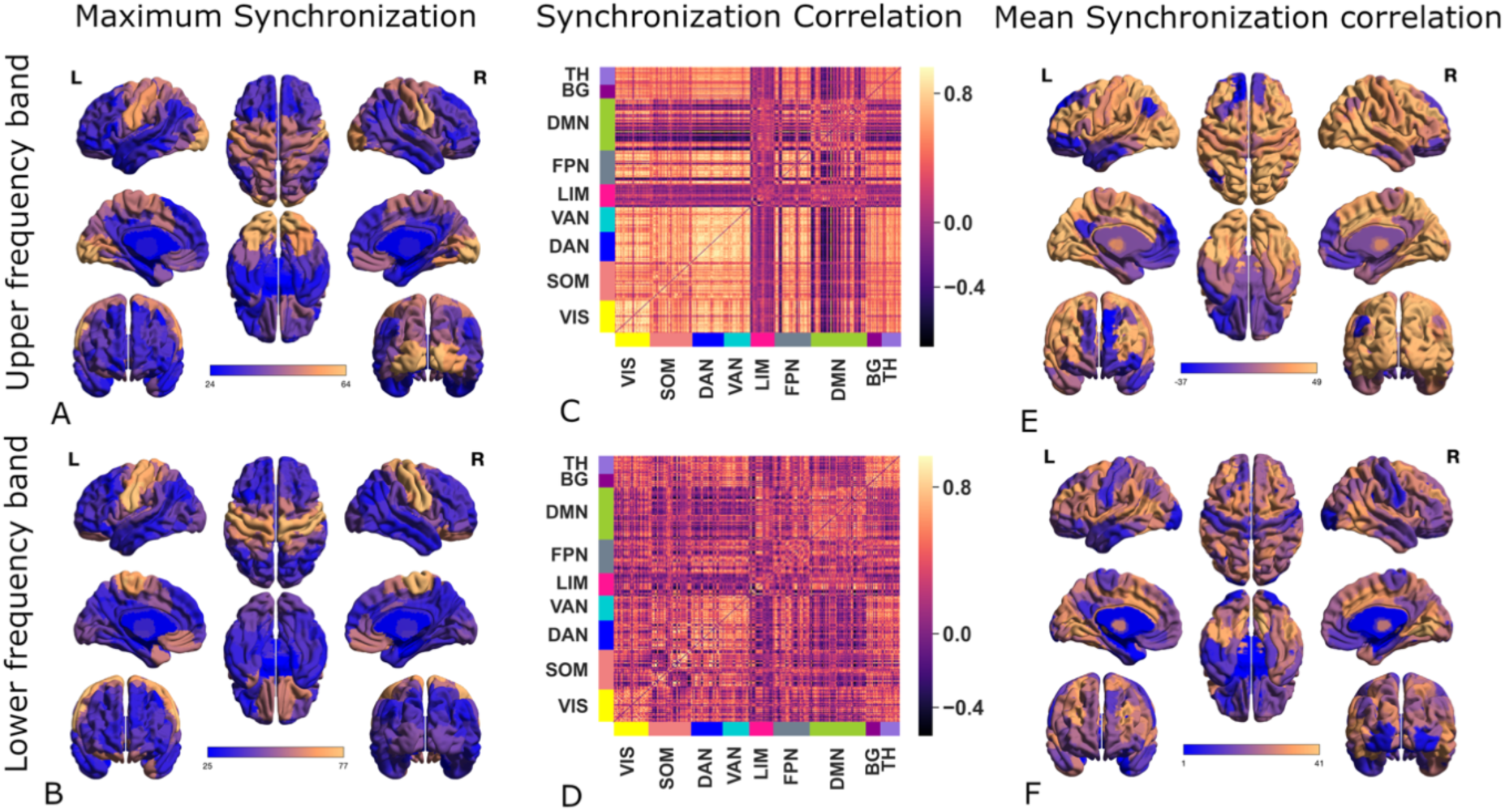
Panels **A** (upper frequency band) and **B** (lower frequency band) show the maximum degree of cross-subject synchronization across time and communities for all the parcels (282 subjects). The network (with left hemispheric parcels initially grouped, followed by right hemispheric parcels) and parcel pair-specific presentations of the mean correlation of community synchronization (averaged separately for COM1-6 and all subjects) over time are shown in panels **C** and **D**, respectively. In panels **E** and **F**, the mean (i.e., by averaging across rows of the correlation matrices shown in panels **C** and **D**) correlation of synchronization for all parcels (Pearson’s r × 100) is shown. The amygdala, hippocampus, and basal ganglia parcels are not shown.

In the upper-frequency band (Fig. 5C), the DMN and limbic networks were decoupled (more so in the left hemisphere than in the right), whereas most parcels in the remaining networks were highly correlated in terms of cross-subject synchronization. An inspection of the mean pattern of synchronization per parcel (that is, by taking the mean of the rows in Figs. 5C and 5D) revealed a distinct pattern of integration (high average correlation) between the VIS, DAN, VAN, Thal, BG, DMN (right hemisphere) networks, and most of the FPN network for the upper frequency band (Fig. 5E). Additionally, limbic and predominantly left-hemispheric DMN regions were segregated in relation to other networks.

In the lower frequency band, the cross-subject synchronization time series for the VIS, attentional (VAN, DAN), FPN, Thal, and BG networks exhibited relatively high intra-network correlations (Fig. 5D). Additionally, the between-network correlation for the Thal and BG networks was strong. Moreover, the SOM network displayed interhemispheric decoupling (Fig. 4), except for the tongue motor areas, which were strongly synchronized (Fig. 5D). The motor and somatosensory cortices, as well as portions of the DMN and visual poles, appeared decoupled from the rest of the brain, as measured by the average correlation of the cross-subject synchronization time series (Figs. 5D, F). The maximum degree of cross-subject phase synchronization exhibited a near-oscillatory pattern, with three peaks per block (Suppl. Fig. S5B).

### Activity of the SMA and cingulo-opercular action mode network

It is well known from the literature that the supplementary motor cortex (SMA), pre-SMA, anterior mid-cingulate cortex (aMCC), insula, supramarginal gyrus, putamen, and thalamus are activated before the motor cortex ^4,18^. In instances where movement is self-paced or coupled with a cognitively demanding task, the prefrontal cortex is also engaged ^2,6^. These regions collectively constitute the cingulo-opercular action mode network (CO-AMN), which encompasses the middle frontoparietal operculum (inferior precentral sulcus), middle precentral sulcus, supramarginal gyrus, anterior prefrontal cortex (aPFC), putamen, and thalamus^19^. This network is also known as a timing network in different contexts^20^.

Our analysis revealed a distinct pattern of cross-subject phase synchronization in the bilateral SMA, right precentral sulcus, left inferior precentral sulcus, and right supramarginal gyrus (SMG) (Fig. 6A, See also Suppl. Figs. S9-10 for regions exhibiting the most pronounced synchronized negative shift across subjects). At the onset of the motor task paradigm and at the conclusion of the two fixation blocks, which were interspersed with the task blocks at irregular intervals, significant peaks in cross-subject phase synchronization (i.e., COM5 and COM6) were observed (Fig. 6B). These corresponded to peaks in the negative shifts in the averaged intrinsic mode signal time series (lower-frequency band) (Fig. 6C). Notably, these negative shifts were also present in the intrinsic mode signal time series of the individual subjects for each fixation period (Fig. 6D). Importantly, the negative shifts in the intrinsic mode signals of the individual subejcts had their equivalent in the raw BOLD time series (Fig. 6E), albeit with a temporal shift, such that the maximum negative deflection was observed slightly before that of the raw BOLD time series (Fig. 6D-E, see also Suppl. Fig. S11). The results were very similar when investigating a subcohort (N = 70) with minimal head movement (Supp. Fig. S12).

**Fig. 6.**
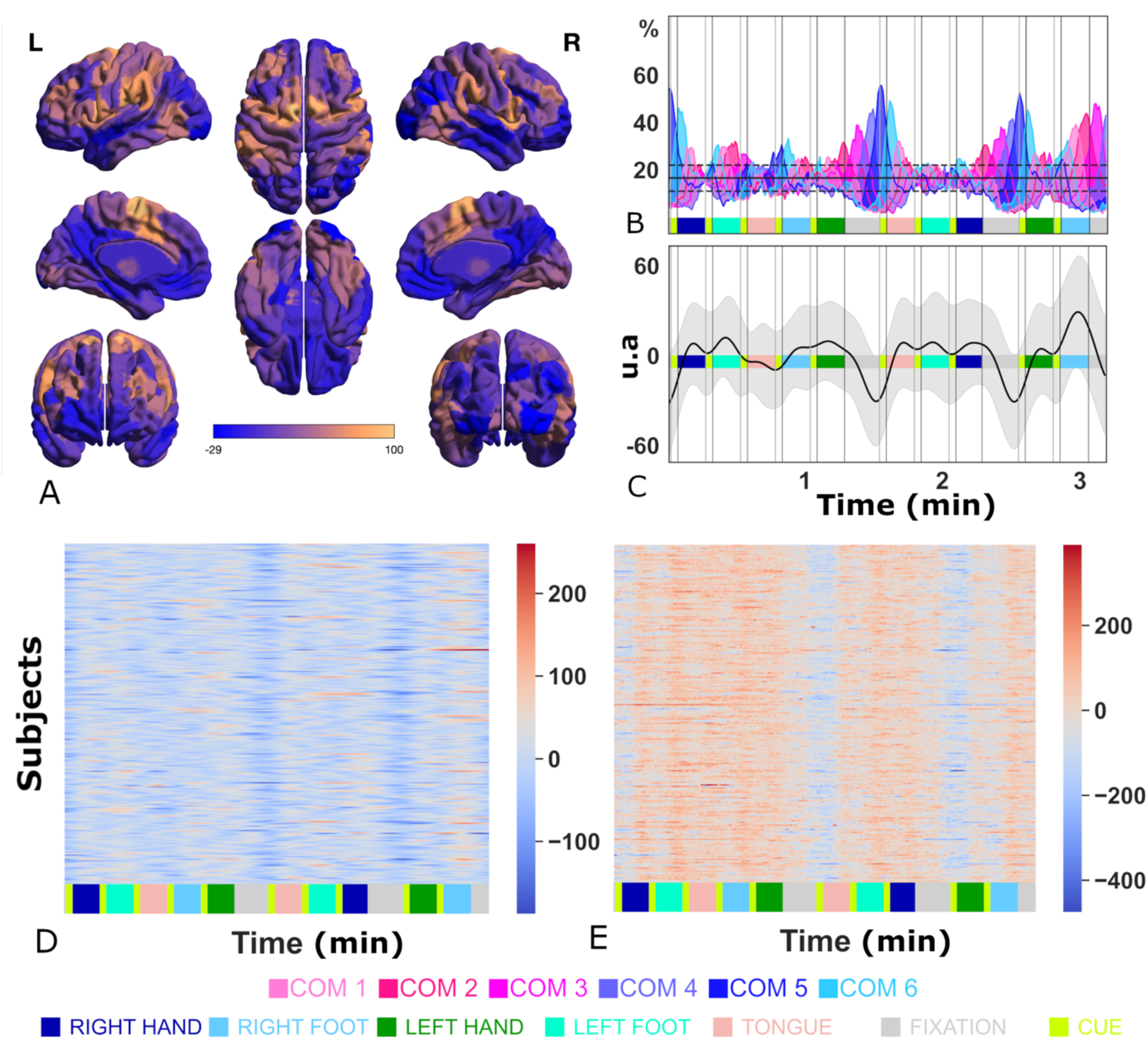
Cross-subject phase synchronization and subject-specific intrinsic mode signals for the left supplementary motor area (LH_SalVentAttn_Med_3) during the fMRI motor task. Panel **A** depicts the mean degree of correlation of cross-subject phase synchronization between the left hemisphere SMA parcel and all other brain regions (Pearson’s r × 100), showing substantial overlap with the CO-AMN network^19^. Panel **B** shows the cross-subject phase synchronization time series (COM1-C; black line represents the mean of the results based on phase-scrambled data, dotted lines = ± 2 SD). The average intrinsic mode signal time-course (lower frequency band) is shown in panel **C**, shaded area = ± 1SD) and the subject-specific time series are shown in panel **D**. For comparison, the raw BOLD signal intensity time-courses are shown in panel **E**.

### Cross-subject synchronization and inter-effector regions

Emerging evidence indicates that the motor cortex can be divided into effector and inter-effector (IE) regions^21^. The effector regions are responsible for the execution of motor movement, whereas the IE regions have been proposed to facilitate whole-body integration and motor planning, displaying non-preferential activation during motor movements. The IE regions exhibit extensive connectivity with each other and with CO-AMN. In the parcellation utilized in this study^17^, only the middle and inferior IE regions were identifiable; specifically, one parcel lateral to the tongue and one situated between the tongue and hand motor regions. Consistent with the literature, we observed that the cross-subject synchronization time series of the IEs were generally highly correlated with each another (Pearson’s r range [0.59, 0.87]) (Suppl. Fig. S13). Contrary to the findings of the Gordon study^21^, which examined individual subjects, our results indicated that IE regions exhibit a preference for specific motor activities. The cross-subject synchronization time series demonstrated a strong positive average correlation with tongue movement (r = 0.75, SD = 0.07), positive average correlation with hand movements (r = 0.49, SD = 0.19), and negative average correlation with foot movements (r = -0.28, SD = 0.12) (average calculated across the four effector regions and two motor regions). There was a stronger average correlation between the IE regions and the SMA network (r = 0.59, SD = 0.10) compared to the average correlation between the IE regions and the rest of the brain (r = 0.18, SD = 0.30).

## Discussion

We employed a new time-resolved fMRI method to demonstrate that the equivalent of the readiness potential is present in a substantial sample of Human Connectome fMRI motor task data^15,16^. During rest (fixation) intervals between motor tasks, a slow and increasingly negative shift in the intrinsic mode BOLD signal was observed bilaterally in the SMA prior to a of forthcoming motor actions within a cued paradigm (block design). Although the negative shift was most pronounced in the SMA, it was also observed in regions largely overlapping with the cingulo-opercular action mode network (CO-AMN) (Fig. 6A, Suppl. Fig. S9), and disappeared after the cue for the upcoming task was presented. CO-AMN is associated with heightened arousal, externally focused attention, cue-specific processing, planning, and processing of interoceptive signals^19^. The SMA is part of the ventral attention network for the parcellation schema used in this study, further attesting to its involvement in attentional processes ^17^. Importantly also, SMA and anterior insula are consistently involved in both reward and loss anticipation in monetary incentive delay tasks^22^. Regions in the CO-AMN have also been shown to be activated by processes involving timing in the brain; therefore, they are referred to as the brain timing network^20,23^. A large meta-analysis of timing studies using fMRI showed that a right hemispheric dominant version of CO-AMN consistently activates during timing tasks involving either perceptual or motor components, or both^23^. This network has a topology very similar to that of the SMA - centred network, which synchronizes during the fixation periods in our results (Suppl. Fig. S9). The minor differences observed in the network boundaries between the CO-AMN, the timing network, and the SMA-centred network reported here are likely a reflection of the flexible engagement of these regions during different task demands, mirroring real-world complexity, which recruits similar, partly overlapping networks. For example, the PFC is not engaged in the motor tasks, which can be attributed to the basic paradigm that lacks significant cognitive complexity ^19^.

CO-AMN has also been found to be strongly correlated with the recently described IE regions^21^. We found similar, but more attenuated, correlations (Suppl. Fig. S13). This discrepancy could be explained by methodological differences between precision mapping in a smaller number of densely sampled individuals and the parcellated large sample used here. Task-specific activation of some IE regions observed in our study is likely to have the same cause.

Importantly, the negative signal shift in the CO-AMN (Fig. 6) was present in single subjects and for single epochs, with some minor differences in timing. These results suggest that the negative shift serves as a general indicator of active anticipation within the CO-AMN for an upcoming motor task. It is also likely to include some degree of motor planning. However, because the exact nature of the upcoming task is unknown during the fixation period (which limb, which side), this planning must be general in scope; thus, no lateralization is seen. Recently, results from an electrophysiological study showed that the RP amplitude increased as a function of learning the optimal time to move. No increase was observed when the timing was random, and learning was impossible. The authors concluded that the RP reflects expectations more strongly than uncertainty ^24^. Our results support the notion that learning the timing of motor movements is reflected in the activity of the SMA and CO-AMN when motor movements are predictable. During the first of the two back-to-back fMRI motor runs (Suppl. Fig. S14), fixation periods were interspersed with three consecutive task blocks. A peak in the negative shift in cross-subject synchronization was observed towards the end of the fixation blocks. It was also observed where the third fixation block would have been placed if it had followed the same order between tasks, which it did not (see Online Methods and Suppl. Fig S17). In the second run, there were no signs of this type of expectation (Fig. 6B). This finding suggests a rapid learning effect (within one run), that fixation periods in this paradigm occurred at irregular intervals. The rapid learning of timing is well documented, as is the involvement of the SMA in timing ^20,23^. In addition, in both runs, the SMA showed a clear negative shift right at the start of the task runs in the same way as at the end of the fixation period. In both runs, there was a positive shift in synchronization at the end of the last task block. We interpret this as a relaxation response, as opposed to a negative shift, indicating anticipation.

While the cross-subject phase-synchronization algorithm closely captured both positive and negative shifts in the intrinsic mode signals, in some instances, the negative shifts were an artifact of the intrinsic mode decomposition, with no clear correspondence in the raw BOLD signal. An example is the pre-cue negative shift in motor movement-specific parcels (Fig. 4, Suppl. Fig. S6). Importantly however, the negative shifts in the intrinsic mode signals of the SMA and CO-AMN during fixation periods had a clear equivalent in the raw BOLD time series, albeit slightly shifted in time (Fig. 6, Suppl. Fig. S9-12).

Our results partially reconcile previous interpretations of the RP in so far that they demonstrate that RP is a genuine phenomenon, observable not only in electrophysiological recordings but also in fMRI data, and detectable at the level of individual subjects and single epochs (Fig. 6). Its bilateral distribution suggests that it is indistinguishable from contingent negative variation (CNV) when lateralization of the upcoming motor task is unknown, supporting the hypothesis that early CNV and RP components largely represent the same type of signal manifesting in slightly different contexts^2,11^. However, the design of the task paradigm examined here does not permit fine-tuned discrimination of potential differences in the late components of CNV and RP signals or conclusions regarding its role in the decisional process^1,10^. While the relationship between the RP and fMRI BOLD response has been investigated in a few studies, those studies have not identified the RP per se in the BOLD signal. Instead, they observed BOLD activations (positive deflections) associated with the negative potentials in the EEG ^4,25^.

The negative shift in the sluggish BOLD signal is naturally more protracted in time than the RP and other negative potentials observed in the neurophysiological recordings with better temporal resolution. Importantly, it appeared in the slower of the two main frequency intervals that were the focus of our analysis. This was clearly differentiated from another type of recruitment of the ventral and dorsal attention networks, which together with the visual network, exhibited task-locked synchronization in the higher-frequency band (Fig. 2). It is well known from neurophysiological recordings that periodic sensory stimulation can induce a rhythmic brain response that mirrors the stimulation frequency ^26^. It has been proposed that slow cortical potentials (SCPs) that are recruited ^6,26^. Given the slow nature of both SCPs and fMRI BOLD signals, they are likely to reflect the same or similar processes, albeit with a time lag^11,12^. In contrast to the regular synchronization patterns in the CO-AMN generated by the block design in the upper frequency band (Fig. 2), the slow negative shift in the CO-AMN in the lower frequency domain (Fig. 6) appears to be driven by an active anticipation process. This means that internally generated and externally presented information is processed in different frequency bands in the same region. This is in line with previous fMRI studies, which have shown differentiated processing in different frequency bands^27–29^. Our results suggest that this processing is sequential and not simultaneous for the SMA, because the peaks in cross-subject synchronization in the lower frequency band are accompanied by desynchronization in the upper frequency band (Suppl. Fig. S15). Finally, we noted previously unreported complex co-activation of the orbitofrontal cortex (OFC), ventromedial prefrontal cortex (vmPFC), and temporal poles for tongue movements involving both positive and negative shifts in the BOLD signal (Suppl. Figs. S7-8).

Taken together, the results of our time-resolved analysis of BOLD signal time courses, based on their instantaneous phases and the corresponding phase-ruled community membership, contribute new knowledge on the role of negative shifts in fMRI signals. Negative BOLD signals are conventionally interpreted as “deactivations”, hence “not active.” Given the results presented here, this notion seems too simplistic^30^. Instead, in analogy with the closely related phenomenon of SCPs, that likely reflect cyclic modulations in overall cortical excitability ^31^, negative shifts in the BOLD-signal does not necessarily correspond to a passive, “non-use” process, but rather an active process of inhibition that significantly impacts the brains’ state-specific responsiveness and it is amenable to subjective intention ^30–33^. De-activation responses, notably deactivation of the DMN during task paradigms, were noted early in the era of fMRI research ^30,34,35^. These anti-correlations between network activity were by some initially thought to be artifacts^36^. More recently, whole-brain approaches have begun to harness information present in the entire brain simultaneously, moving away from the bias of activation versus deactivation ^32,37–40^. Our new method demonstrated cross-subject synchronization through cycles of both positive and negative shifts in the BOLD signal, further highlighting the importance of the whole-brain perspective. It also provides a methodological approach for investigating anticipation and attention in a wide variety of fMRI paradigms. Given the importance of anticipation in perception and the creation of subjective reality, as well as its close association with prediction, it provides an exciting opportunity to investigate these phenomena further ^41–44^.

## Material and methods

### Data

fMRI data from 381 young and healthy participants of both sexes from the HCP 1200 subject release were initially included in this study ^15,45^. The two motor fMRI task runs (block design, scanned back-to-back) used here were acquired at the end of a close to an one-hour fMRI recording session^16^. In the first run (run A), phase-encoding was in the right-left direction and the second (run B) phase encoding was in the left-right direction. Subjects with head movement with a peak frame-wise displacement root mean square (fdrms) of > 0.5 mm were excluded from each of the two runs (See Fig. 1 for fdrms time series, run B). In the first run, 266 subjects remained, and in the second run, 262 subjects remained. Because the cues of most motor tasks were associated with very minor average head movements in this group, we also performed the same analysis on a subset of subjects with very low motion (max fdrms <0.2mm) (Suppl. Fig. S16).

The second run (run B) was the focus of the main analysis. When appropriate, to highlight important differences, results from the first run (run A) will also be included. The motor task consisted of five movements (finger-tapping left, finger-tapping right, toe squeezing right, toe squeezing left, and tongue movement)^16^. The duration of each block was 12 s and was preceded by a 3 second cue. Each type of movement/block was repeated once. Additionally, three fixation blocks (duration: 15 s) were interspersed between motor task blocks. The ordering of movements in runs A and B was different, but some of the basic structure was preserved (Suppl. Fig. S17).

### Preprocessing and parcellation

Minimally pre-processed volume-based data provided by the HCP were used; no further pre-processing of the data was conducted^46^. The pre-processing pipeline included gradient unwarping, motion correction in six directions, distortion correction using a field map, nonlinear registration to the MNI reference space using the FSL tool FNIRT, and intensity normalization^16,46^. For cortical parcellation, the Schaefer 200 brain area parcellation (seven networks) was used^17^. It is important to note that the parcels in the Schaefer parcellation are not symmetrical across hemispheres. Furthermore, the subcortical section of the Human Brainnetome Atlas^47^ was used to delineate the thalamus, basal ganglia, hippocampus, and amygdala. In total, the parcellation comprised of 236 brain parcels. The amygdala and hippocampus parcels were assigned to the “limbic network” in the Schaefer parcellation scheme, whereas the thalamus and basal ganglia were treated as separate networks. The analysis therefore considered nine separate networks: Visual (VIS), Somato-motor (SOM), Dorsal attention (DAN), Ventral attention (VAN), Limbic (Limb), Frontoparietal (FPN), Default Mode (DMN), Thalamus (Thal), Basal Ganglia (BG). This was the same parcellation used in previous studies ^39,40^.

### Complex weights, instantaneous phase, and Complete Ensemble Empirical mode decomposition (CEEMD)

Time series analysis is an example in which complex numbers are ubiquitous. A real signal S can be represented in analytical (i.e. complex-valued) form using polar coordinates *S* = *A* × *e*^*iθ*^, where A is the amplitude of the signal, theta is the angle and *i* the imaginary unit. The same signal can be expressed in Cartesian coordinates as *S* = *A*(cos(*θ*) + *i* × *sin*(*θ*). The cosine term represents the real part of the phase angle and the sine component represents the imaginary part.

Signal phase is one of the classic measures used to assess co-activation between regions of the brain ^48,49^. It has mostly been used in MEG and EEG studies and to a lesser extent in fMRI data analysis ^39,40,49–52^. Phase synchronization is the alignment of the phases of two signals. This can refer to the relationship between two signal time series. In the idealized case, the phase difference between the two time series is constant over time. It can also refer to a single time point by estimating the instantaneous phase using the Hilbert transformation^53^. The assumption is that a small difference in phase translates into information exchange between regions^48^. Thus, the smaller the difference, the stronger the cooperation. An important methodological strength of phase as a measure of interregional relationships in the brain is that it quantifies relationships on a continuous spectrum of positive and negative values. This is conceptually similar to correlation, which also spans positive and negative dimensions. The instantaneous phase provides the added advantage of being time-resolved. Previous studies have shown high similarity between time-resolved changes in connectivity between BOLD-signals based on correlation and instantaneous phase measures respectively^39,54,55^. Importantly, the instantaneous phase extracted using Hilbert transform must be unambiguous. This implies that the time series cannot contain riding waves, which are typically present in real-world signals. Narrow-band pass filtering is the most common method for making a time series smooth with unambiguous phase representation at each time point. The frequency range of the BOLD-signal that contain neuronally associated information remains an open question. Some argue for a very narrow range, while there is growing evidence pointing to a wide range of meaningful frequencies within the BOLD-spectrum^51^. In fact, there seems to be task-specific segregation of network activation in different frequency bands, which is also present to some extent during rest ^27,28,56–58^. Thus, a narrow filter is likely resulting in information loss, which may be relevant. Also, the choice of frequency cut-off is often arbitrary.

Considering these factors, we opted to use an adaption of the empirical mode decomposition (EMD) algorithm called Complete Ensemble Empirical Mode Decomposition (CEEMD). The EMD is a heuristic sifting algorithm that decomposes a signal into oscillatory modes called intrinsic mode functions ^59^. In each mode the total number of maxima and zero crossings differ by at the most one. This means that there will be no riding waves and no ambiguity regarding phase. The waveforms are locally approximately symmetric ^59^. This, by design, forces a negative deflection prior to and after a positive deflection (albeit it could be very minor) whether it is present in the raw signal or not. Because the combined modes reconstruct the original signal, an imposed deflection is compensated in the subsequent modes. To interpret the results correctly, it is necessary to examine the original time series. To reduce the downstream effects of this process, we first applied a broad bandpass filter ([0.003,0.3] Hz) to reduce the noise because the (CE)EMD extracts the highest frequencies first. This also substantially reduced mode mixing. The CEEMD is a noise-assisted development of the EMD which results in a complete decomposition of the signal with only a small error and improved separation of frequencies, that is, reduced mode mixing ^60^. Due to edge effects, 10 time points at the beginning and end of each time series were removed.

### Phase clustering algorithm

As a starting point, we define the dense complex weighted matrix *M*_*t*_ where *t* is the time. Dense here means that all pairwise relationships between parcels are included without threshold. Each entry *M*_*t*_(*x*, *y*) is an edge that describes the instantaneous phase difference *θ*_*x*,*y*_ between parcel (node) *x* and *y* at time *t*. Complex weighted networks are also an emerging research topic in the network science community, and we refer interested readers to, for example, Böttcher and Porter 2024 and Tian and Lambiotte 2024, for more theoretical support^61,62^. This algorithm is greedy and has a hierarchical structure. It optimizes two main criteria: integration (criterion one) and segregation (criterion two). Integration in this context means that *all* pairwise relationships within the same community have a maximum phase difference defined by threshold one, *thr*1 = cos (*θ*_1_). We investigated six different choices for 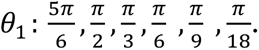 In this context, an integrated community refers to a community that exhibits *phase consistency*, meaning that the maximum phase difference between any pair of nodes within the community is less than *θ*_1_. Segregated means that *all* pairwise relationships between nodes in two different communities have a *minimum phase di=erence* defined by threshold two, *thr*2 = cos (*θ*_2_), where *θ*_2_ = −*θ*_1_. There are also two secondary, more generous thresholds, to account for scenarios where lesser degrees of segregation are present: *thr*2*b* = cos (*θ*_2*b*_), where 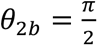 and *thr*2*c* = *thr*1. *Thr*2*b* is only relevant when *θ*_1_ < *θ*_2*b*_ or in situations where no cluster is formed based on *thr*2. *Thr*2*c* is applied only when segregation occurred within this range [*θ*_1_, *θ*_2*b*_]. Thresholds beyond *thr*2*c* are superfluous because it equals a scenario when there is only a single phase-consistent cluster based on *thr*1. In each iteration, the algorithm aims to identify a pair of maximally segregated (criterion two), phase-consistent (integrated) communities (criterion one) if a minimum segregation of *at least Thr*2*c* (= *thr*1) exists in the data. In each iteration, to harmonize communities across time points, the first community is based on the parcel with the smallest instantaneous phase value among the unassigned parcels, that is - *π* in the idealized case. The second community is defined based on the parcel most segregated from the seed parcel of the first community. The first community is subsequently expanded by finding groups of parcels that fulfil both criteria one and two. If multiple such groups exist, the group with the smallest mean phase difference with the seed parcel is chosen. The second community was expanded in the same manner. Communities formed beyond the first iteration were sorted based on their leading and lagging relations with the communities in the first iteration. The imaginary component (*sin)* of the phase difference between two communities provides information on which community is leading, and which is lagging. Let 𝖯_1_ be the instantaneous phase of *Com1* and 𝖯_2_ that of *ComX*. Then, if sin(𝖯_*x*_ − 𝖯_1_) > 0, *ComX* is leading in relation to *Com1*. However, this is a truth with modification since 𝖯_1_ = 𝖯_1_ + 2*π* in which case Com1 would be ahead of *ComX*. If a leading community already exists, a secondary leading community is formed. If instead sin(𝖯_*x*_ − 𝖯_1_) < 0, *Com X* lags in relation to *Com1*.

The results for 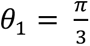 are presented in the main text. The algorithm was designed for community comparison between multiple iterations of a complex weighted network, where the distribution of phases was assumed to be similar across iterations. In the case of time series, the iterations are discrete time points. The algorithm can be adapted to situations in which the phase distribution varies widely between the iterations. This is done using the iteration with the largest distribution as the reference.

### Phase shuffled data

To test whether the results were not merely driven by chance, the algorithm was applied to surrogate data. Because the key property of the investigation was phase, phase shuffled data were used as surrogates^63^. The shuffling of the phases preserves the power spectrum of a time course while destroying the temporal order of the phase. The intrinsic mode time series of the second run (run B) was shuffled. Thus, the shuffled data provided statistical thresholds for significant results. Because of edge effects resulting from the phase shuffling process, 30 time points were excluded at the beginning and end of each time series, instead of only 10, which were excluded from the real data.

## Supplement

**Suppl. Fig. S1.**
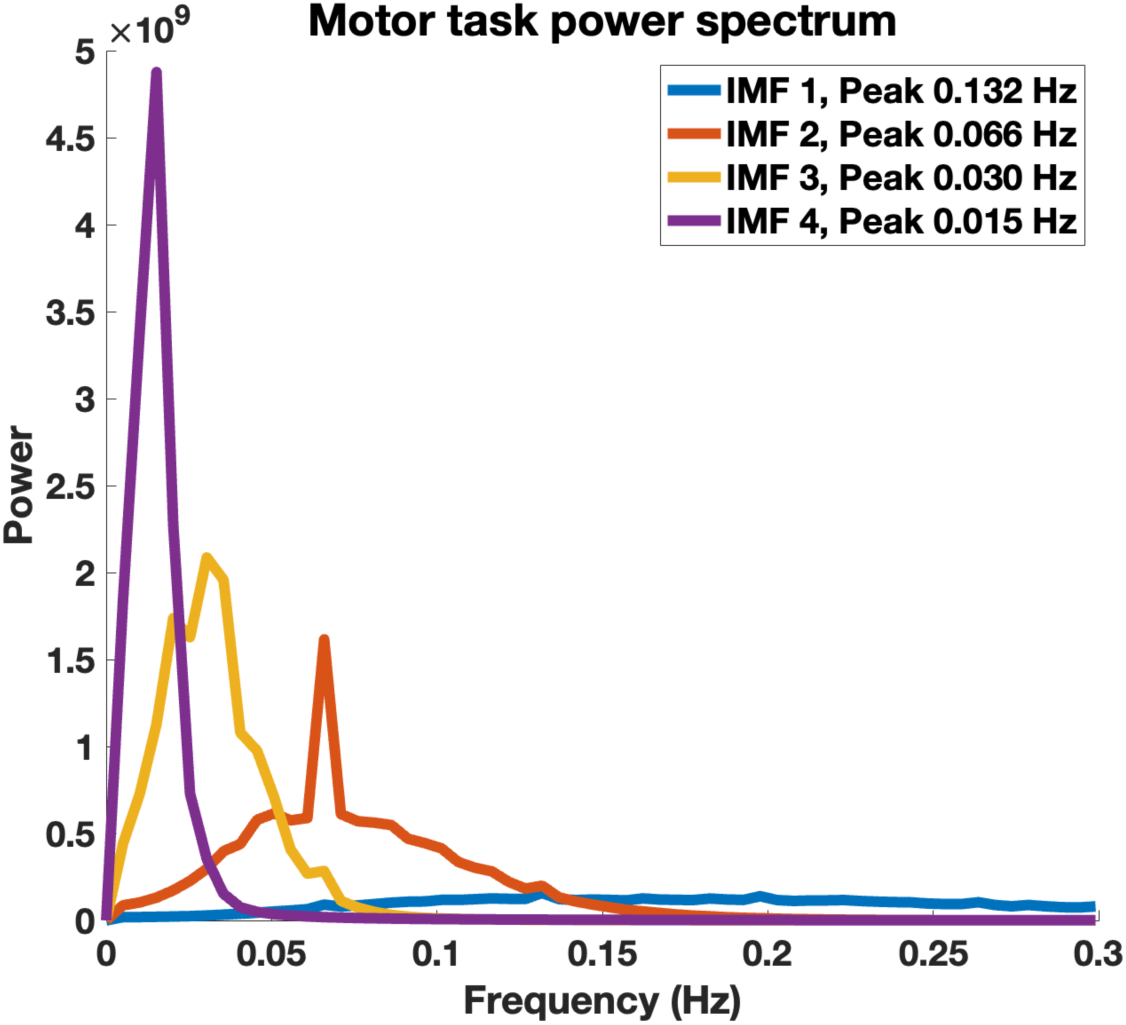
Power-spectrum for four intrinsic mode functions (IMFs) (final run). IMF1 [0.1318, 0.4613] Hz, **IMF2 (upper frequency band) [ 0.0C53, 0.187C] Hz**, IMF 3 **(lower frequency band) [0.0304, 0.03C3] Hz**, IMF4 [0.0152, 0.1064] Hz. Only results from upper and lower frequency bands are presented.

**Suppl. Fig. S2.**
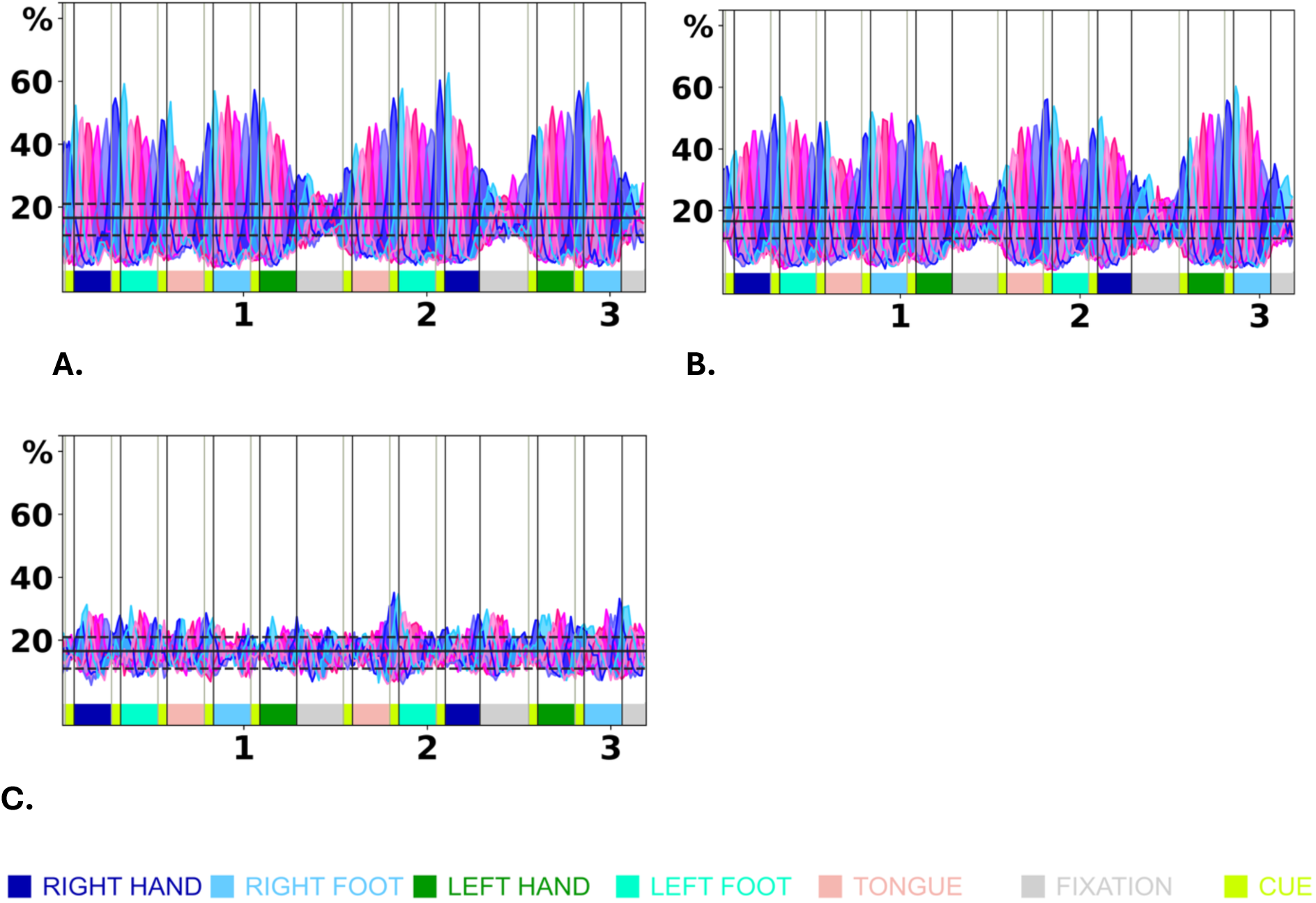
**A)** and **B)** example of a visual network parcels cross-subject synchronization, upper frequency band desynchronizes during fixation blocks (grey), second run. **C)** Visual parcel that doesn’t show significant cross-subject synchronization. X-axis is time in minutes. Y-axis is percent of total subjects.

**Suppl. Fig. S3.**
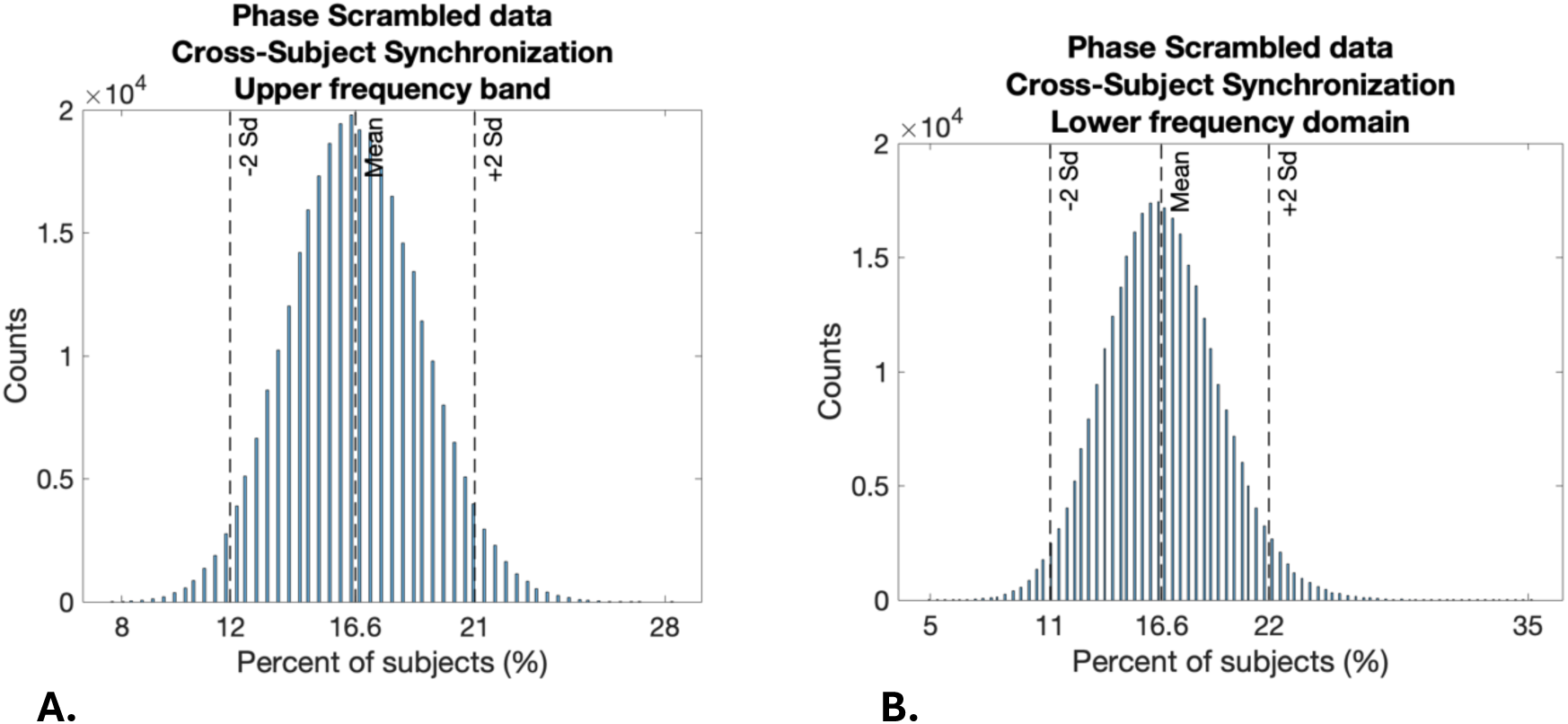
Distribution of cross-subject synchronization across all six communities, time-points and subjects (count) based on the phase scrambled data. Due to edge e=ects of the phase scrambling 15 timepoints were cropped at the start and finish. **A)** upper frequency domain, **B)** lower frequency band.

**Suppl. Fig. S4.**
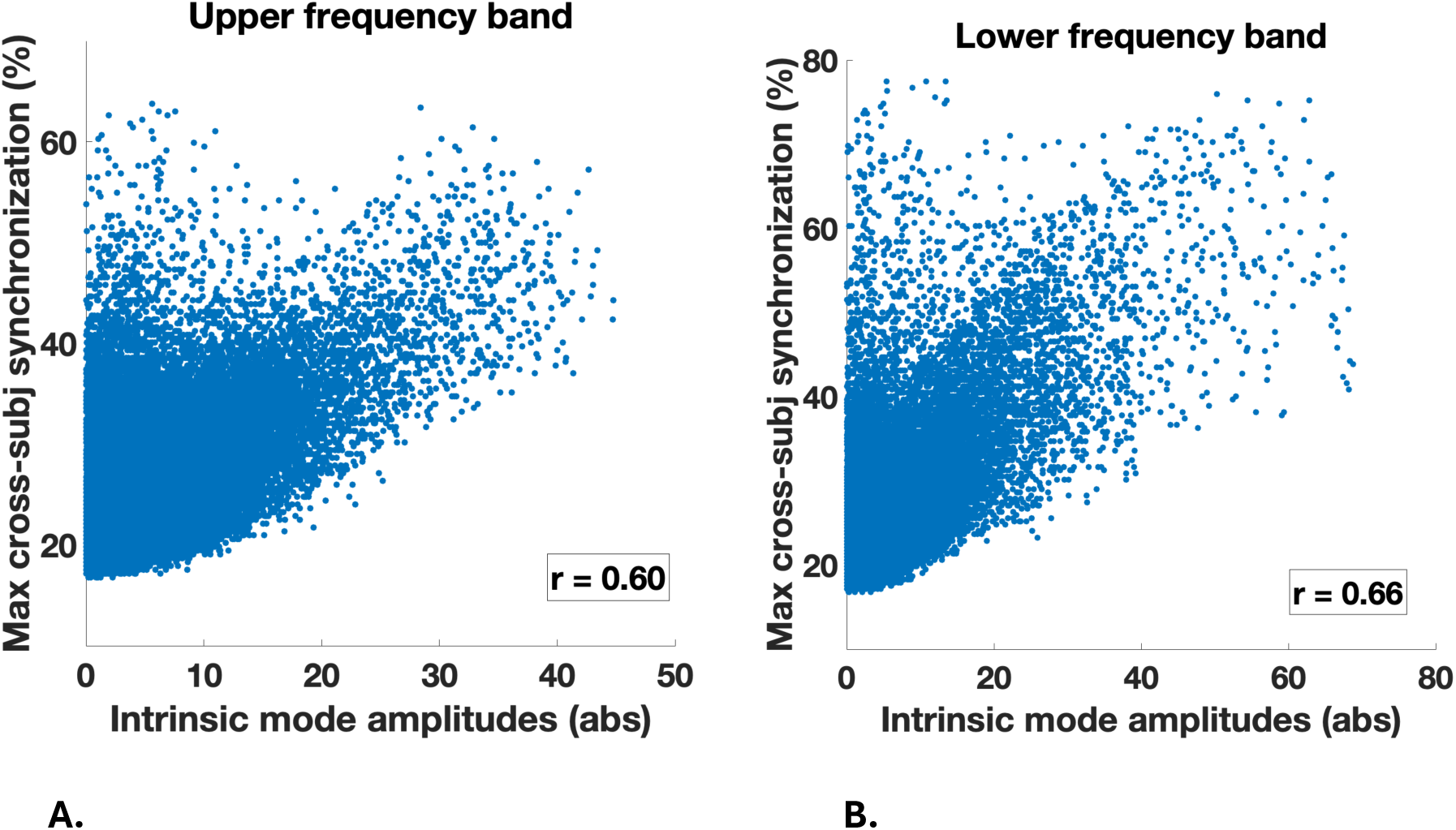
Scatterplots showing the relationship between intrinsic mode signal amplitude (absolute value) and maximum (among the six communities) cross subject phase synchronization for all of the 23C parcels and 2C4 time points in the time-series. **A)** Upper frequency band, Pearsons’ r = 0.C0, **B)** Lower frequency band, Pearsons’ r = 0.CC.

**Suppl. Fig. S5.**
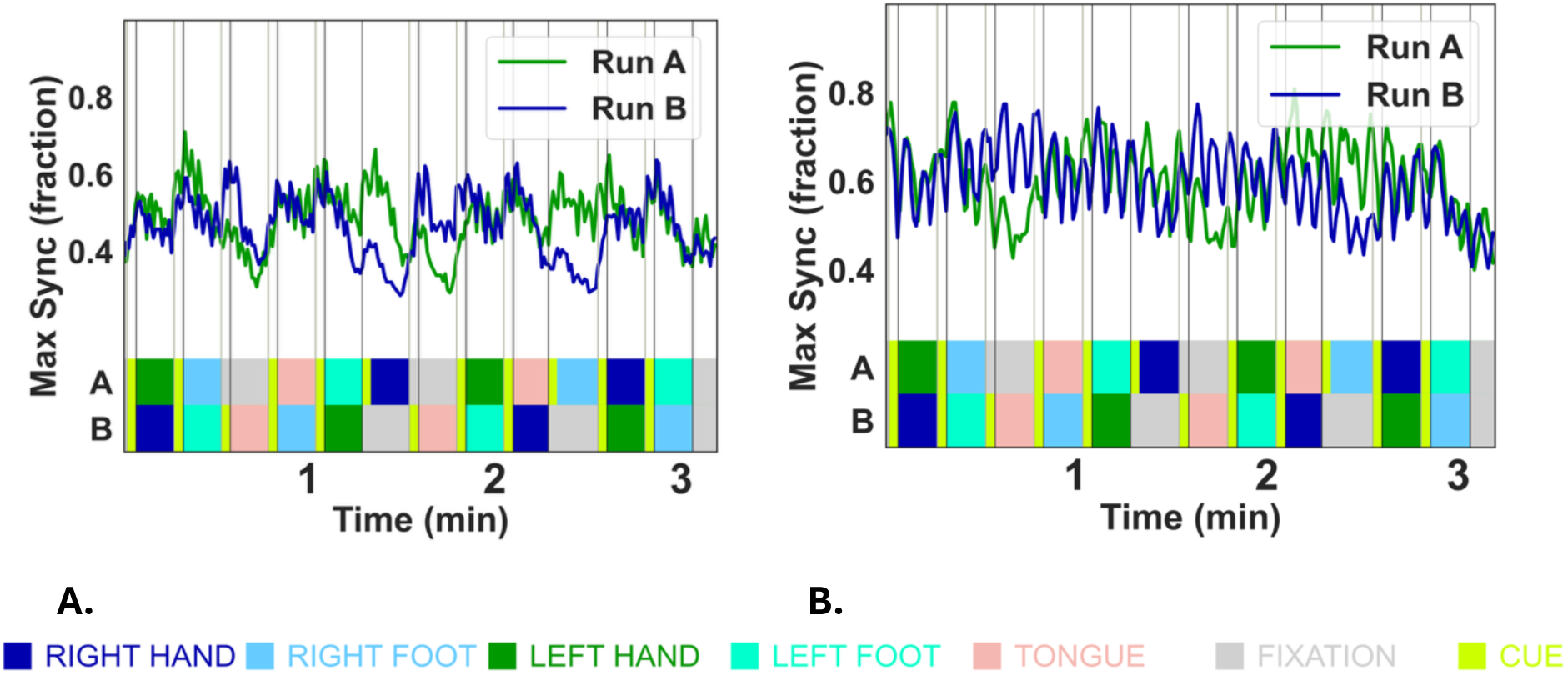
Maximum cross-subject phase synchronization across the six main communities as a function of time in the first (Run A) and second (Run B) run **A)** Upper frequency band, **B)** lower frequency band. Synchronization is generally higher in the lower compared to the upper frequency band. The lower frequency band exhibit a near oscillatory pattern with three peaks per block.

**Suppl. Fig. S6.**
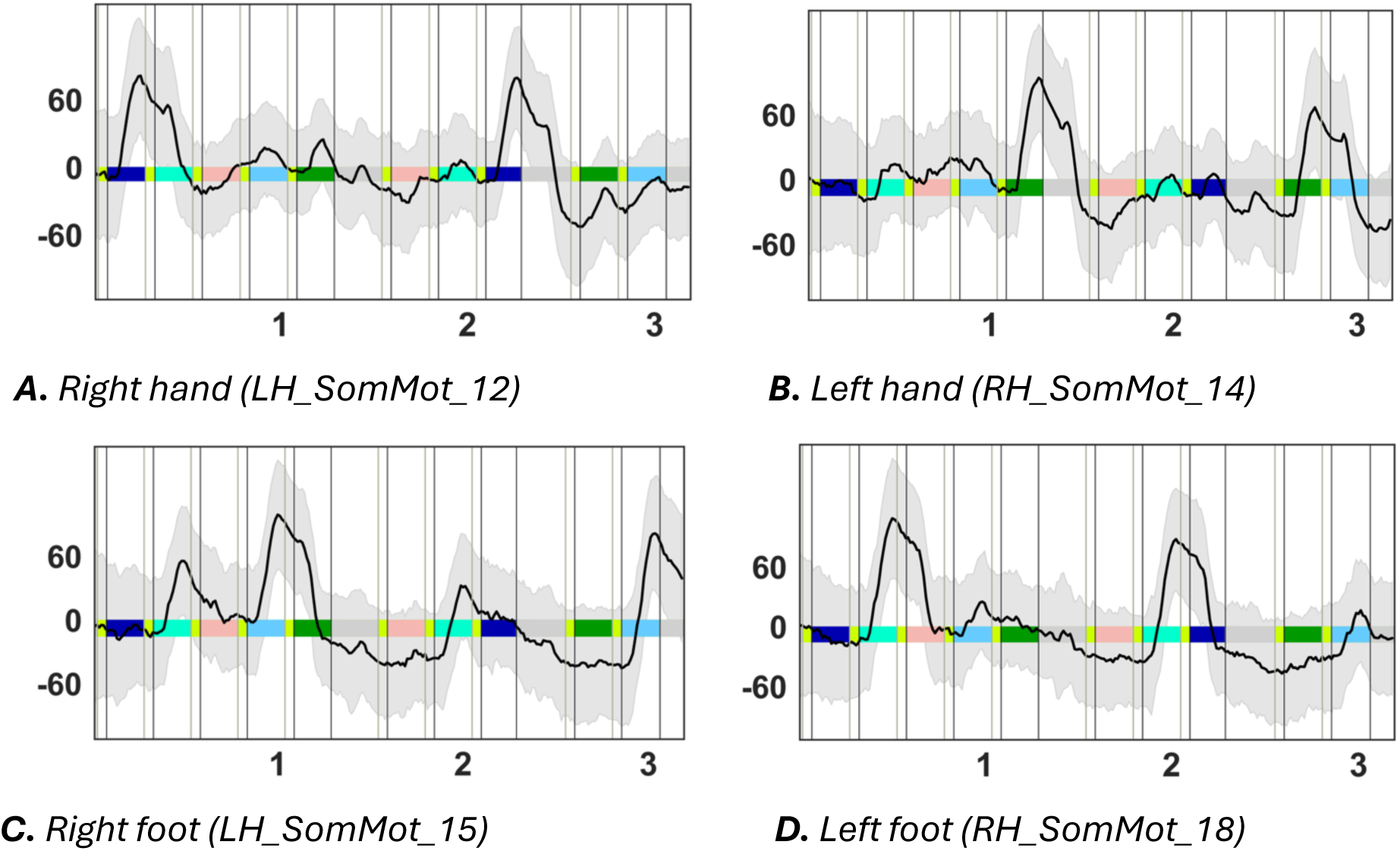
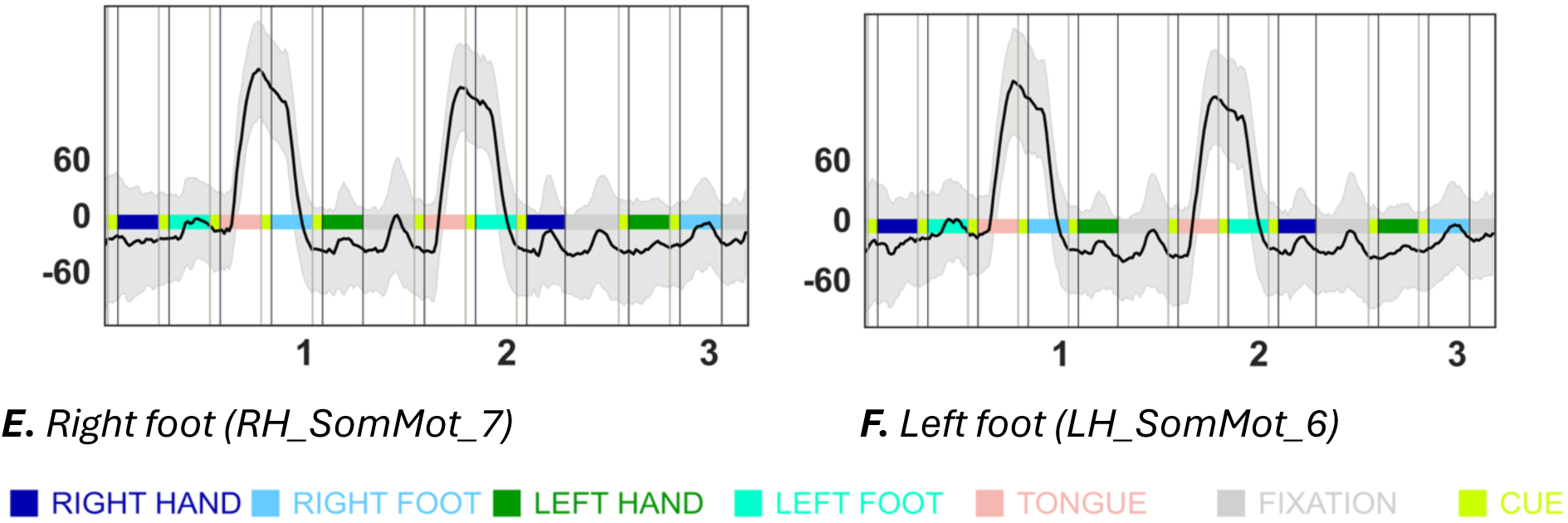
Averaged (n=282) raw BOLD time-series for the six movement specific parcels in the motor cortex (shaded area = ± 1 SD)). Tongue motor movement parcels from both hemisphere motor cortex are included (E, F). Comparing the averaged raw BOLD with the lower frequency intrinsic modes in Fig. 4 suggests that the pronounced negative deflections seen in the latter prior to task execution in this case are artefacts of the CEEMD-algorithm (See Methods). X-axis is time in minutes. Y-axis is arbitrary units.

**Suppl. Fig. S7.**
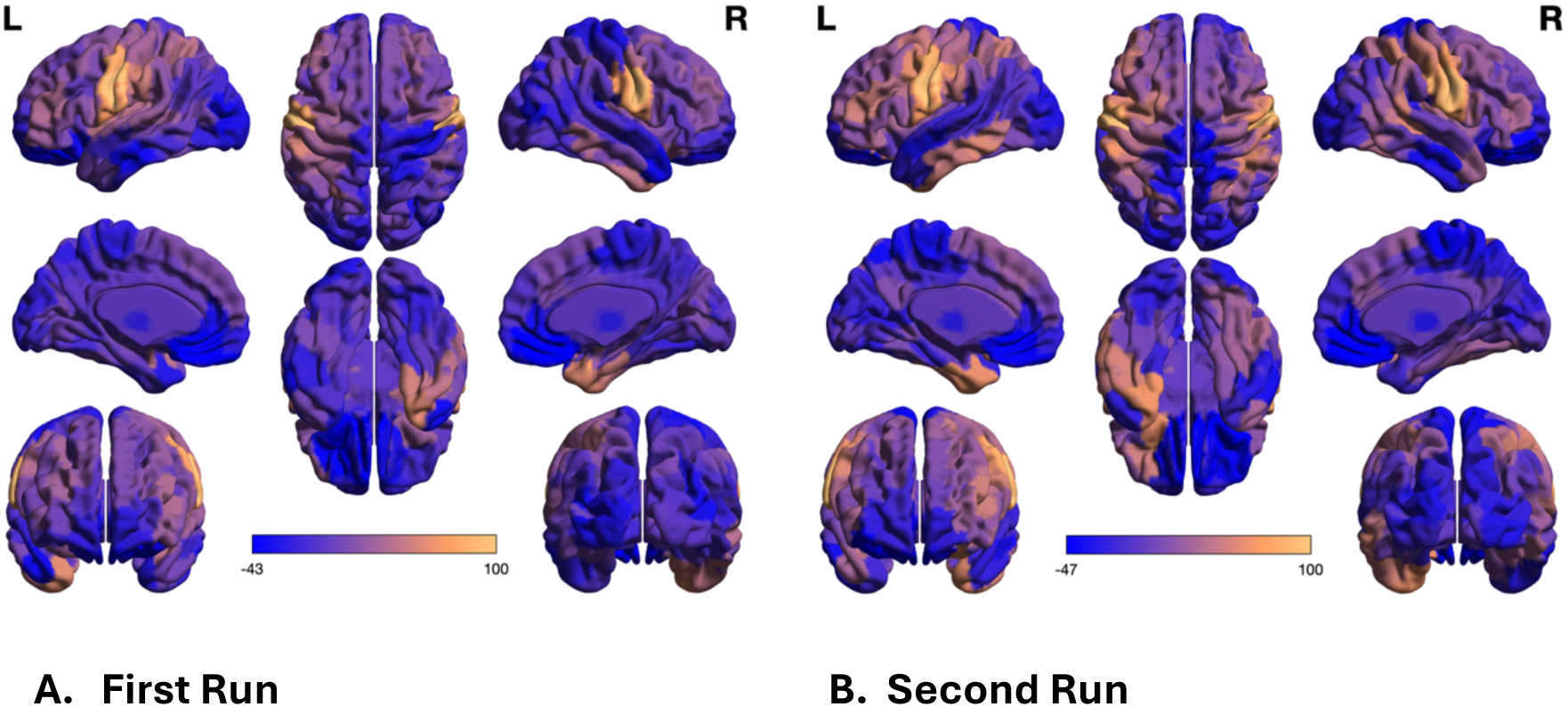
Cross-subject phase synchronization correlation maps for tongue movement, Iower frequency band. A) First Run. B) Second Run (main results). Tongue movements induced synchronization in the temporal pole, orbitofrontal cortex (OFC), ventromedial prefrontal cortex (vmPFC), and specific regions of the basal ganglia, notably in the left ventral caudate nucleus. These synchronizations were antiphase, or segregated, relative to the tongue motor areas, except for the left temporal pole in the second run. During the two experimental runs, tongue movements engaged the cerebral hemispheres in a partially opposite manner. In the initial run, in-phase synchronization was observed in the right temporal pole rather than in the left, and above-baseline synchronization with the basal ganglia was absent. Note the opposite correlation pattern in the ventral temporal lobes and left lateral OFC between the two runs.

**Suppl. Fig. S8.**
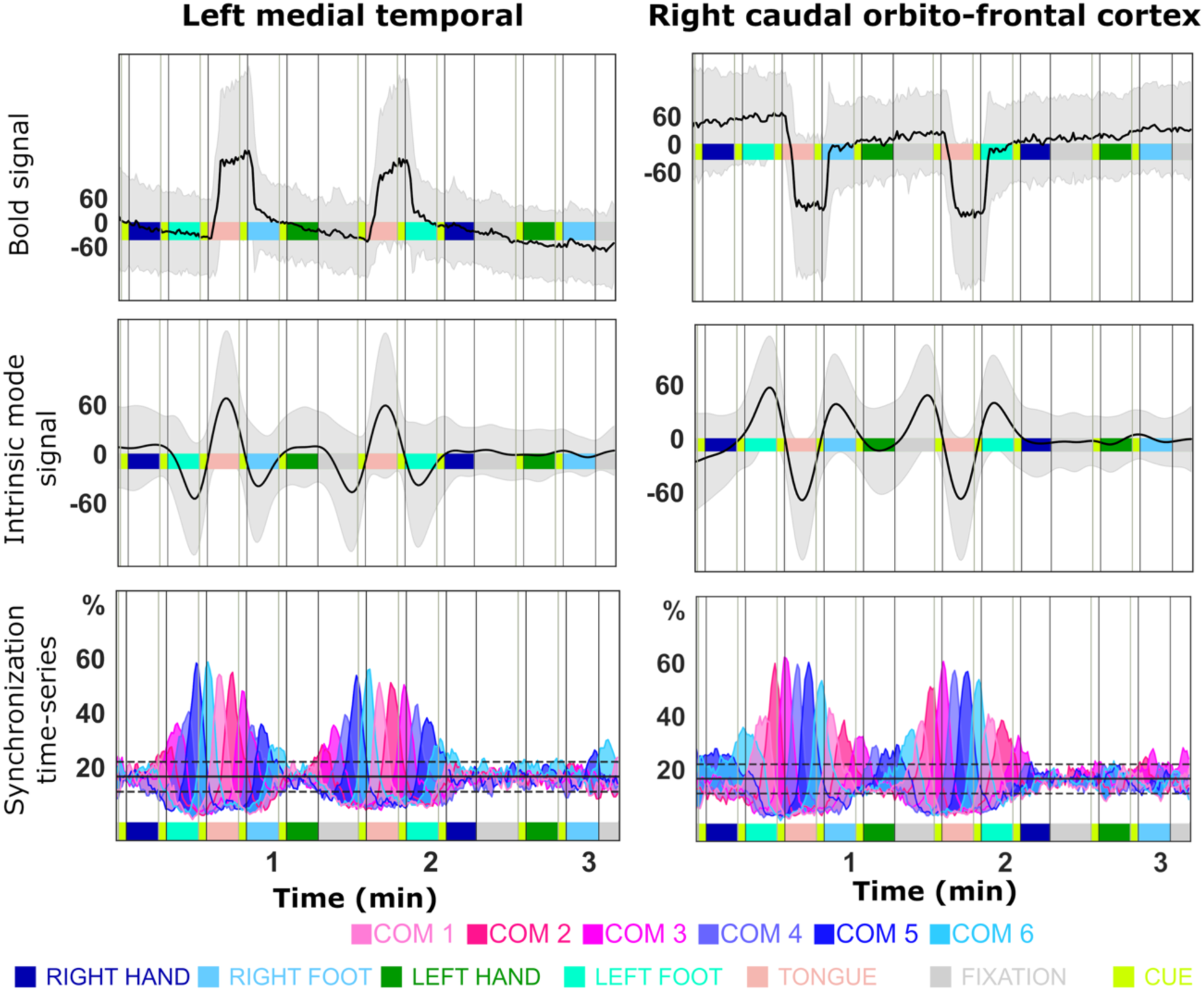
Example from to parcels with opposite signal changes during tongue movement (second run). Left column: Left medial temporal pole (LH_Limbic_TempPole_3). Right column: Right caudal orbito-frontal cortex (RH_Limbic_OFC_2). First row: mean raw BOLD signal (+/- 1 SD in grey). Second row: Mean intrinsic mode signal (+/- 1 SD in grey). Third row: Cross-subject synchronization time-series.

**Suppl. Fig. S9.**
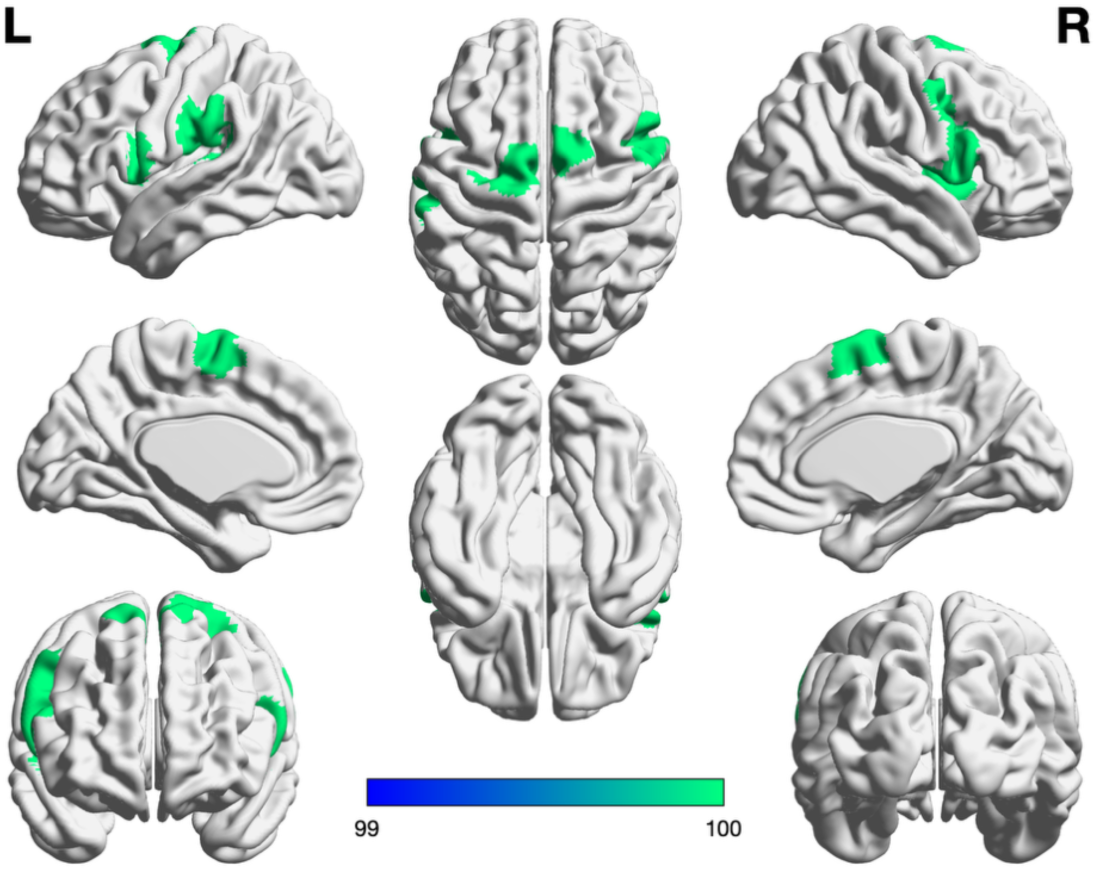
The parcels most strongly connected with the left hemispheric SMA: right SMA, right middle and inferior precentral sulcus, left inferior precentral sulcus and right supramarginal gyrus (SMG). Strong connection was defined as the combination of a) mean correlation of synchronization with SMA r > 0.75 and b) minimum peak in cross-subject synchronization > 40% of total subjects.

**Suppl. Fig. S10.**
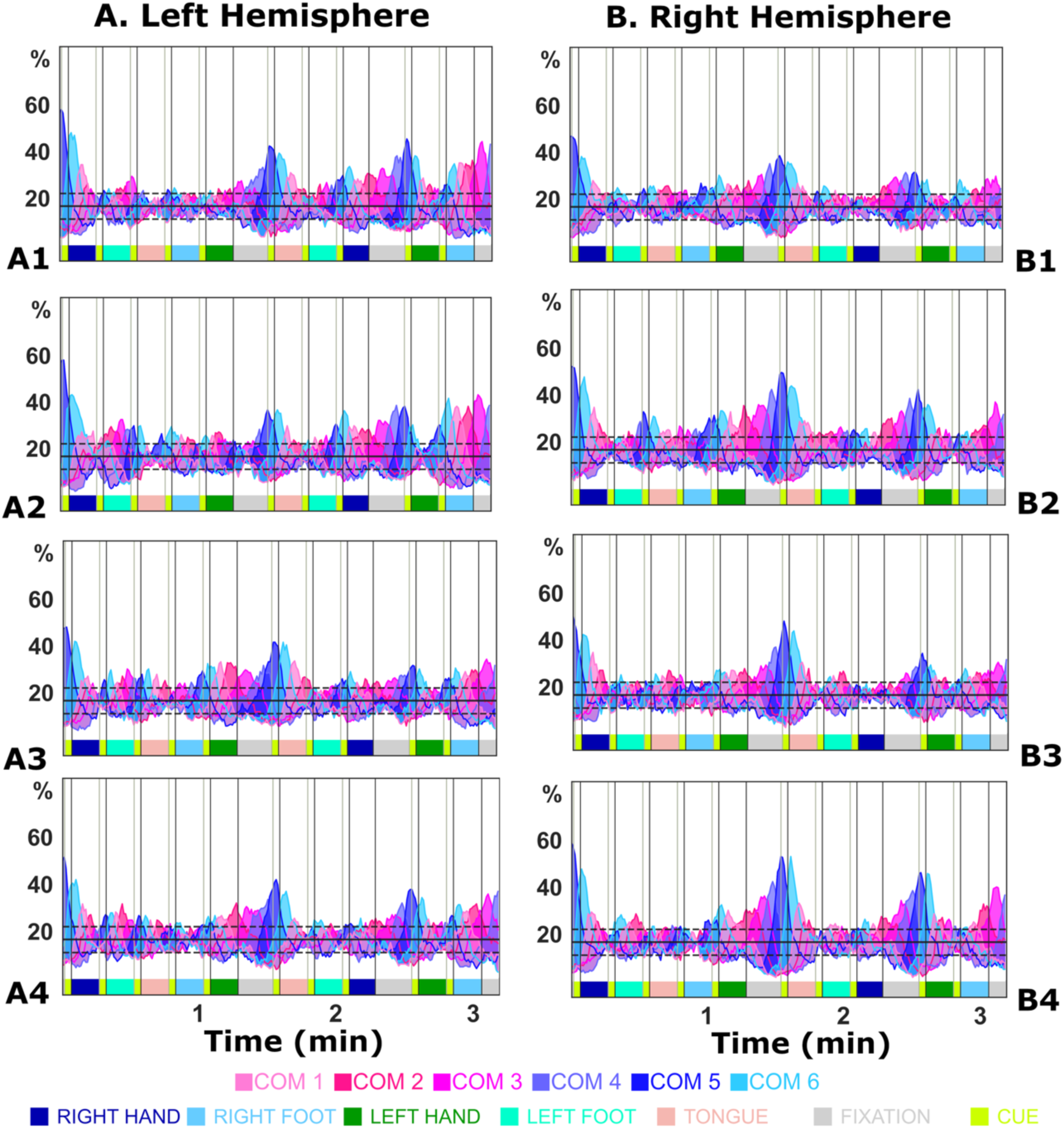
Cross-subject synchronization time-series for the parcels in Suppl. Fig S9, i.e. the parcels with the strongest connection with the left hemispheric SMA. **A1.** LH_SomMot_5, **A2.** LH_SomMot_14, **A3.** LH_SalVentAttn_ParOper_2, **A4.** LH_SalVentAttn_FrOperIns_4, **B1.** RH_SalVentAttn_TempOccPar_1, **B2.** RH_SalVentAttn_PrC_1, **B3.** RH_SalVentAttn_FrOperIns_4, **B4.** RH_SalVentAttn_Med_3.

**Suppl. Fig. S11.**
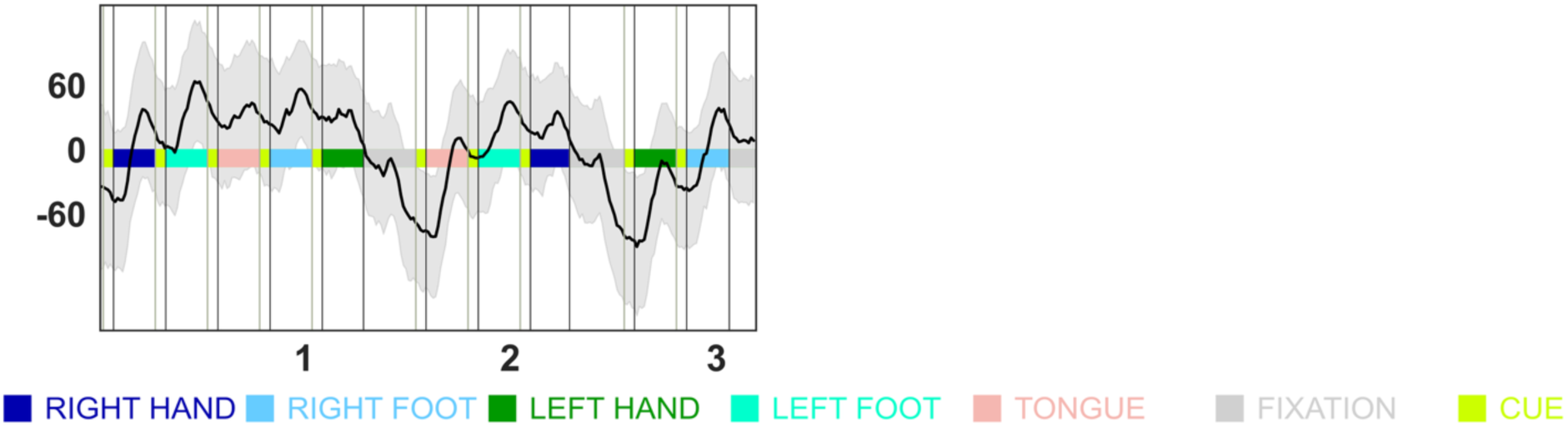
Left SMA (LH_SalVentAttn_Med_3) raw averaged (across subjects) BOLD time-series. X-axis is time in minutes. Y-axis is arbitrary units.

**Suppl. Fig. S12.**
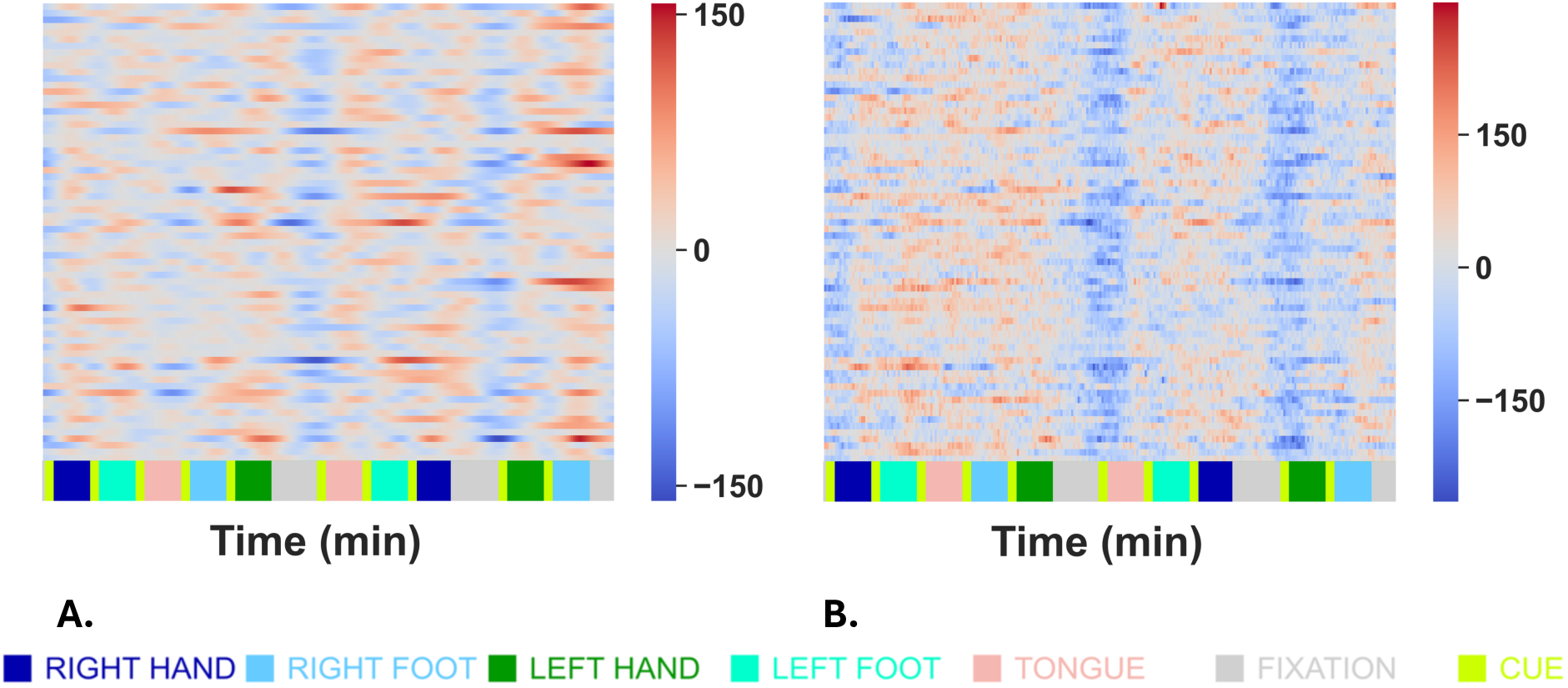
Low motion subjects (N = 70, maximum frame-wise displacement root mean square (fdrms) < 0.2 mm), individual left SMA time series, A) Intrinsic mode time-series lower frequency band, B) Raw BOLD time-series

**Suppl. Fig S13.**
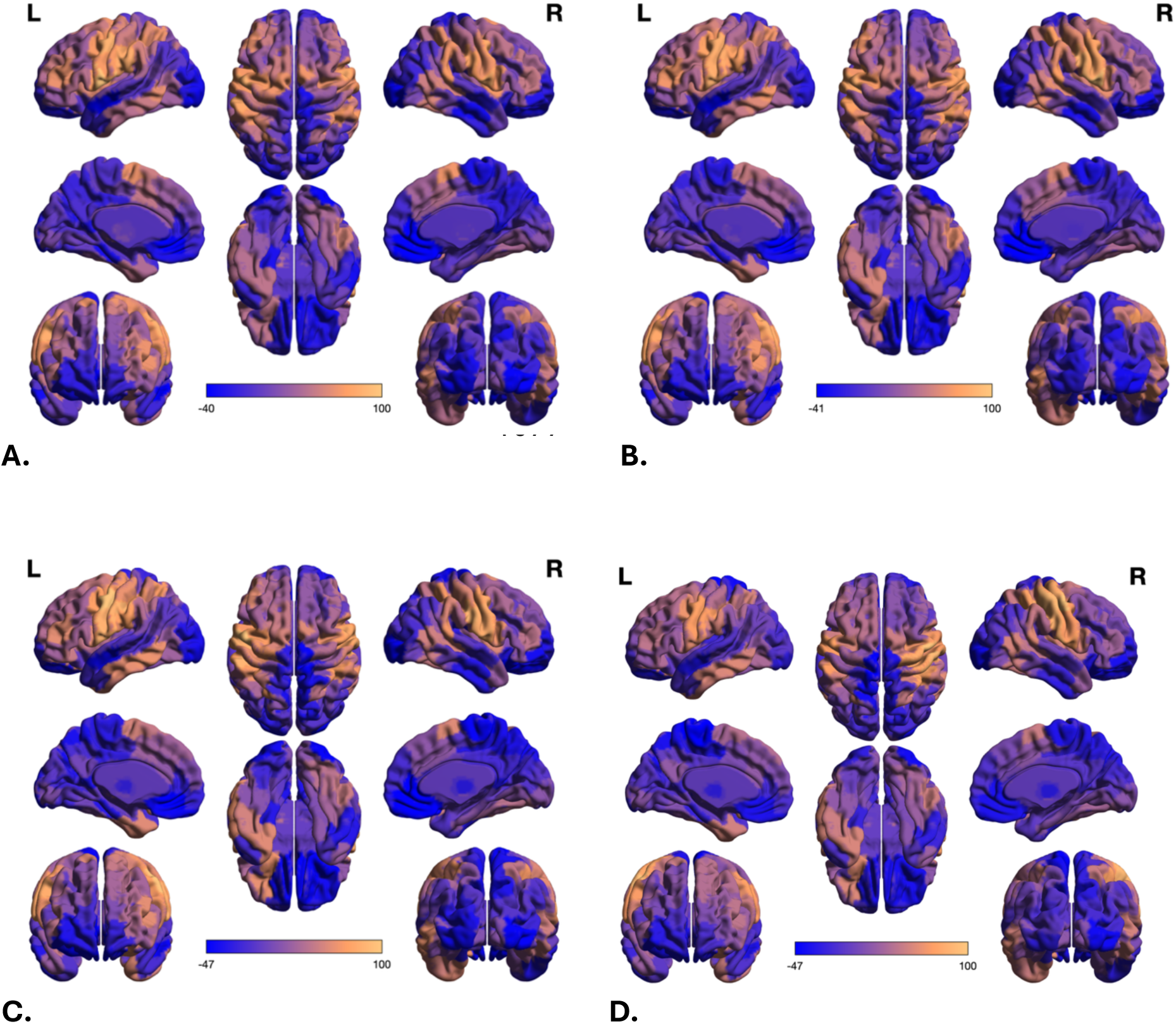
Inter-e=ector regions average correlation of the six coss-subject phase synchronization timeseries with the rest of the brain (Range is Pearson’s correlation x100). A) Left inferior IE, B) Right inferior IE, C) Left middle IE, D) Right middle IE.

**Suppl. Fig. S14.**
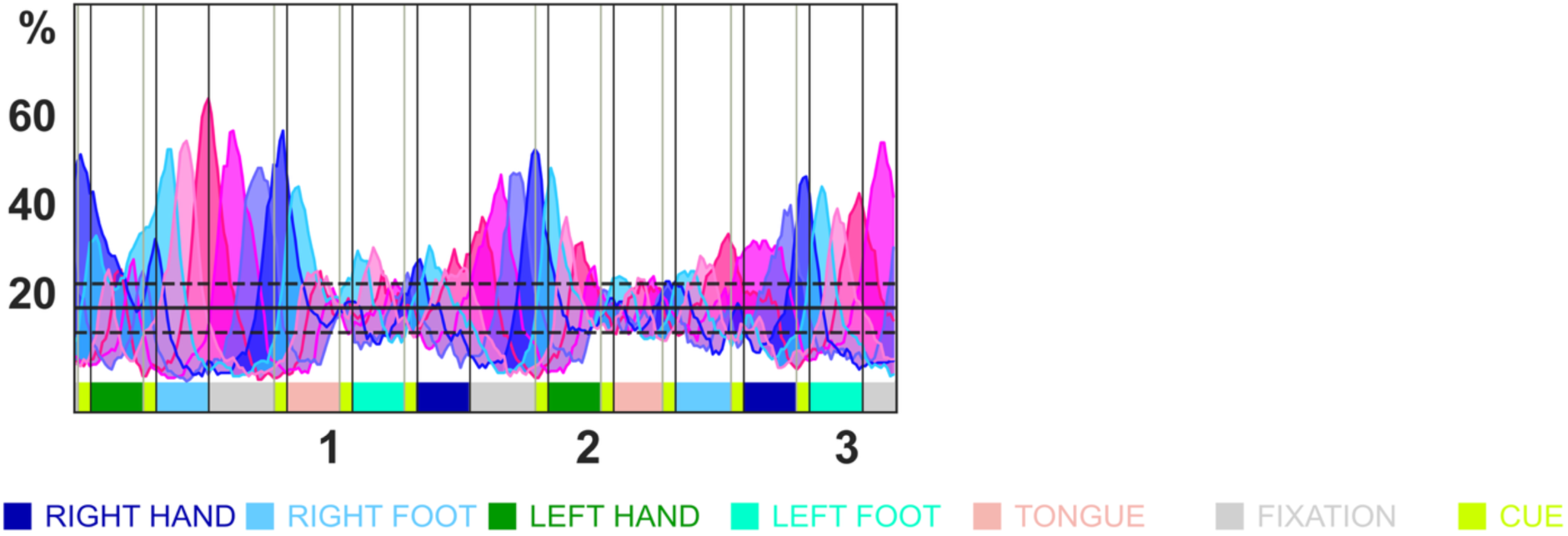
First run (A), Cross-subject phase synchronization time-series for left SMA (LH_SalVentAttn_Med_3). Note the blue peak at the second last task. This is the location the last fixation block would have been following the pattern of the first two fixation blocks. X-axis is time in minutes. Y-axis is percent of total subjects.

**Suppl. Fig. S15.**
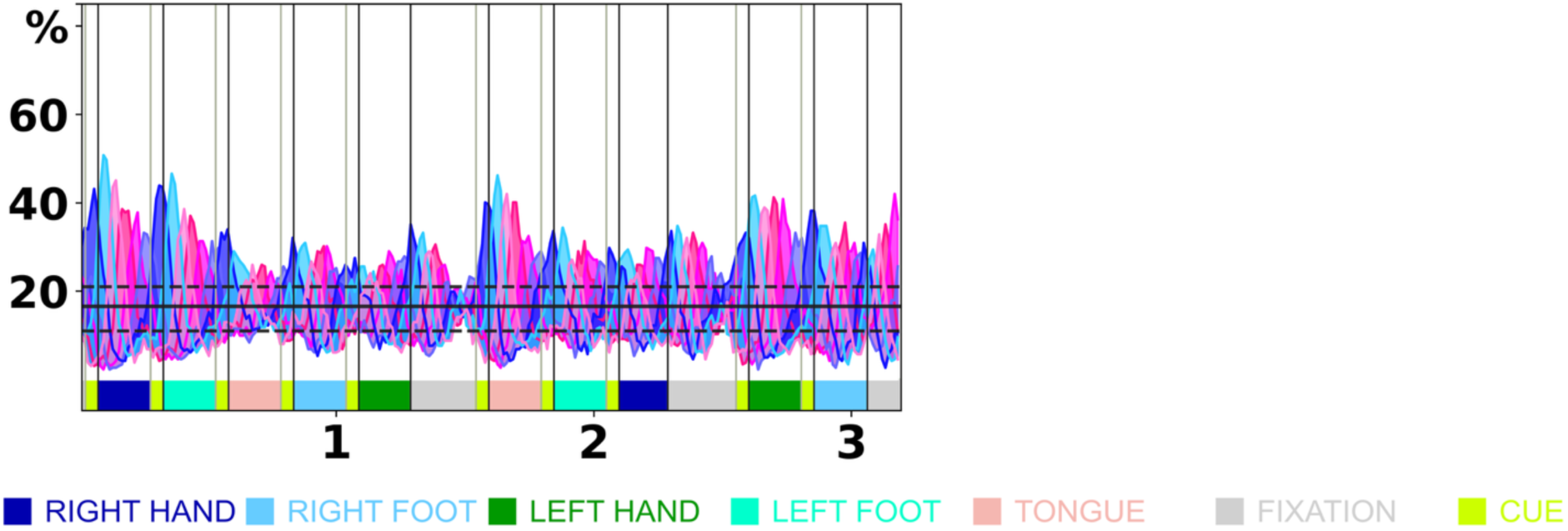
Left SMA (LH_SalVentAttn_Med_3), upper frequency band. Desynchronizations are noted at the end of the two first (i.e. full-length) fixation blocks. The last fixation block is cut due to edge artefacts of the band-pass filtering and intrinsic mode decomposition (see Methods). X-axis is time in minutes. Y-axis is percent of total subjects.

**Suppl. Fig. S16.**
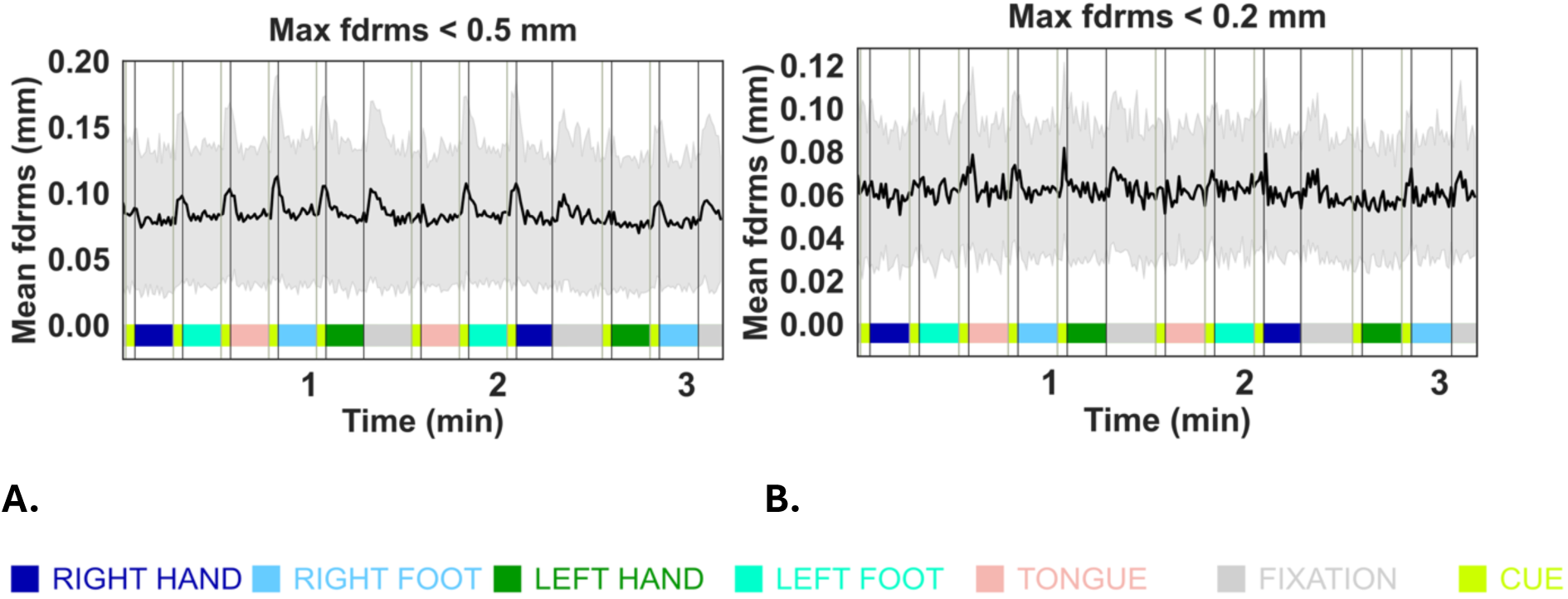
Mean (across subjects) fdrms time series for the second run (run B). A) thr < 0.5mm (N= 262), B) thr < 0.2 mm (N = 70). ± 1 SD in gray shades.

**Suppl. Fig. S17.**
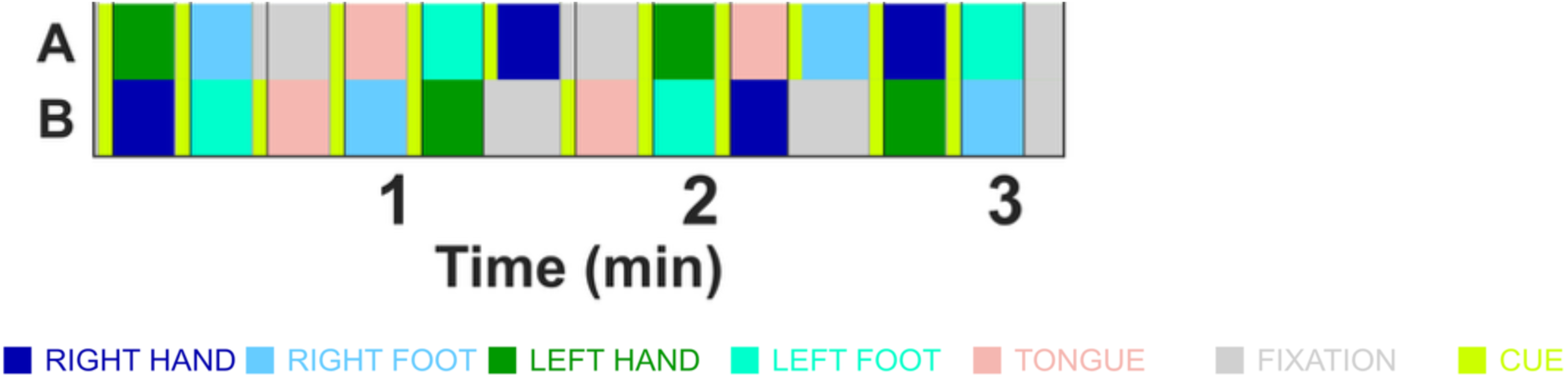
Block design for the two runs: first run (A) and second run (B).

